# Osteoblast-Specific Wnt Secretion is Required for Skeletal Homeostasis and Loading-Induced Bone Formation in Adult Mice

**DOI:** 10.1101/2021.03.30.437737

**Authors:** Lisa Y. Lawson, Michael D. Brodt, Nicole Migotsky, Christopher Chermside-Scabbo, Ramya Palaniappan, Matthew J. Silva

## Abstract

Wnt signaling is critical to many aspects of skeletal regulation, but the importance of Wnt ligands in adult bone homeostasis and the anabolic response to mechanical loading is not well documented. We inhibited Wnt ligand secretion in adult (5-mo) mice using a systemic (drug) and a bone-targeted (genetic) approach, and subjected them to axial tibial loading to induce lamellar bone formation. Mice treated with the porcupine inhibitor WNT974 exhibited a decrease in bone formation in non-loaded limbs as well as a 54% decline in the periosteal bone formation response to tibial loading. Similarly, within 1-2 weeks of Wls deletion in osteoblasts (Osx-CreERT2;Wls^F/F^ mice), skeletal homeostasis was altered with decreased bone formation and increased resorption, and the anabolic response to loading was reduced 65% compared to control (Wls^F/F^). These findings establish a requirement for Wnt ligand secretion by osteoblasts for adult bone homeostasis and the anabolic response to mechanical loading.

## Introduction

Mechanical loading is a potent stimulus for bone formation, mediated in part by the Lrp/Wnt signaling pathway (Robinson et al., 2006). The Wnt co-receptor LRP5 is required for the response to loading (Sawakami et al., 2006), and loading-induced downregulation of the Wnt antagonists Sclerostin *(Sost)* and *Dkk1* favors bone formation (Lara-Castillo et al., 2015; Robling et al., 2008; Tu et al., 2012). While it is well known that Wnt pathway activation is initiated by the binding of one of 19 Wnt ligands to Frizzled and Lrp5/6 cell surface co-receptors (Clevers & Nusse, 2012), the role of Wnt ligands in the anabolic response to loading has not been well described.

Reports by several groups demonstrate that loading induces the expression of Wnt genes in bones of mice. Robinson et al. assayed several Wnt-related genes and reported elevated *Wnt10b* expression 4 h after loading (Robinson et al., 2006). More recently, Kelly et al. used RNAseq and identified *Wnt1, Wnt7b* and *Wnt10b* as genes elevated early after axial tibial loading (Kelly et al., 2016), findings corroborated by others (Galea et al., 2017; Holguin et al., 2016). We recently reported, based on RNAseq, that *Wnt1* and *Wnt7b* were two of only eight transcripts significantly upregulated in cortical bone 4 h after 1, 3 and 5 daily bouts of axial tibial loading in 5-mo old mice (Chermside-Scabbo et al., 2020). Wergedal et al. (Wergedal et al., 2015) reported an increase in *Wnt16* expression with tibial bending in wildtype mice, but a diminished periosteal response to tibial bending in 10-wk old *Wnt16* knockout mice, indicating a functional role of Wnt ligand regulation in the bone response to loading in young mice. (Wergedal et al., 2015)

Wnt protein secretion into the extracellular environment depends on the actions of two intracellular molecules, Porcupine and Wntless, both of which are required for survival in mice (Barrott et al., 2011; Ching & Nusse, 2006; Fu et al., 2009). Porcupine *(Porcn)* catalyzes the addition of a fatty acid to newly synthesized Wnt proteins in the endoplasmic reticulum (Kadowaki et al., 1996; Willert et al., 2003). The modified proteins can then interact with Wntless (*Wls*/GPR177), the Wnt-specific transporter that shuttles vesicle-bound Wnt proteins to the cell surface (Banziger et al., 2006). Inhibition of porcupine using WNT974 blocks bone formation and increases bone resorption in young (8-12 wk old) mice (Funck-Brentano et al., 2018; Madan et al., 2018). Moreover, constitutive deletion of Wntless (*Wls*) in early osteoblasts (Osx-Cre) leads to severely impaired skeletal developmental and perinatal lethality (Osx-Cre) (Tan et al., 2014), while constitutive deletion in mature osteoblasts (Ocn-Cre) leads to osteopenia and spontaneous fractures by 2-mo age (Zhong et al., 2012). Further, results of genome-wide association studies (GWAS) identify 89 variants in the human WLS gene as being significantly associated with bone mineral density (Morris et al., 2019; “Musculoskeletal Knowledge Portal,”). While these studies show a requirement for Wnt ligand secretion by osteoblasts during skeletal development and maturation, it remains unclear whether Wnt secretion by osteoblasts is required for bone homeostasis or the bone anabolic response to mechanical loading.

Our objective was to test the hypothesis that Wnt secretion by osteoblasts is essential for homeostasis in the adult skeleton, and also for the anabolic response to mechanical loading. We dosed adult (5-mo) mice with the porcupine inhibitor WNT974 and observed a decrease in bone formation and increase in resorption, as well as a marked decline in the anabolic response to loading. We then conditionally deleted Wls in osteoblasts in adult mice using a tamoxifen inducible Cre and found that within 1 week, skeletal homeostasis was altered and the anabolic response to loading was greatly impaired. Our findings indicate that secretion of Wnt ligands by osteoblasts is required for adult bone homeostasis and for the anabolic response to mechanical loading.

## Results

### Loading-induced bone formation is reduced in WNT974-treated mice

To investigate the role of Wnt proteins in the anabolic effect of skeletal loading, 5-month old C57BL/6 female mice were subjected to 5 days of cyclic compressive loading on the right tibia while contralateral left tibias served as control. Mice were treated with Porcupine inhibitor WNT974 to block Wnt secretion or vehicle (Fig. 1A). Tibial loading stimulated *periosteal* bone formation in both treatment groups (Fig. 1B); indices of periosteal bone formation were significantly greater in the loaded vs contralateral non-loaded tibias (Fig. 1C-E). However, the periosteal loading response was diminished in WNT974-treated mice. Two-factor ANOVA indicated that loading and treatment had significant effects on Ps.MS/BS, Ps.MAR, and Ps.BFR/BS. Additionally, there was a significant (*p*<0.05) or near-significant (*p*=0.083) loading-treatment interaction for Ps.BFR/BS and Ps.MS/BS, respectively, indicating that the effect of loading depended on treatment. Specifically, in non-loaded tibias, indices of *periosteal* bone formation did not differ between treatment groups. In contrast, in loaded tibias Ps.MS/BS was 41% lower in WNT974-treated mice relative to vehicle-treated mice (*p*<0.001, Fig. 1C), and Ps.MAR and Ps.BFR/BS were 40% (*p*<0.05, Fig. 1D) and 55% (*p*<0.001, Fig. 1E) lower. Similarly, when expressed as a relative measure (i.e., loaded minus non-loaded control), loading-induced periosteal bone formation was reduced by 54% in the WNT974-treated group (Suppl. Fig. S1). Together, these data confirm that loading potently induces periosteal bone formation in the adult skeleton, and indicate that the periosteal loading response depends on the secretion of Wnt proteins.

**Figure 1.**
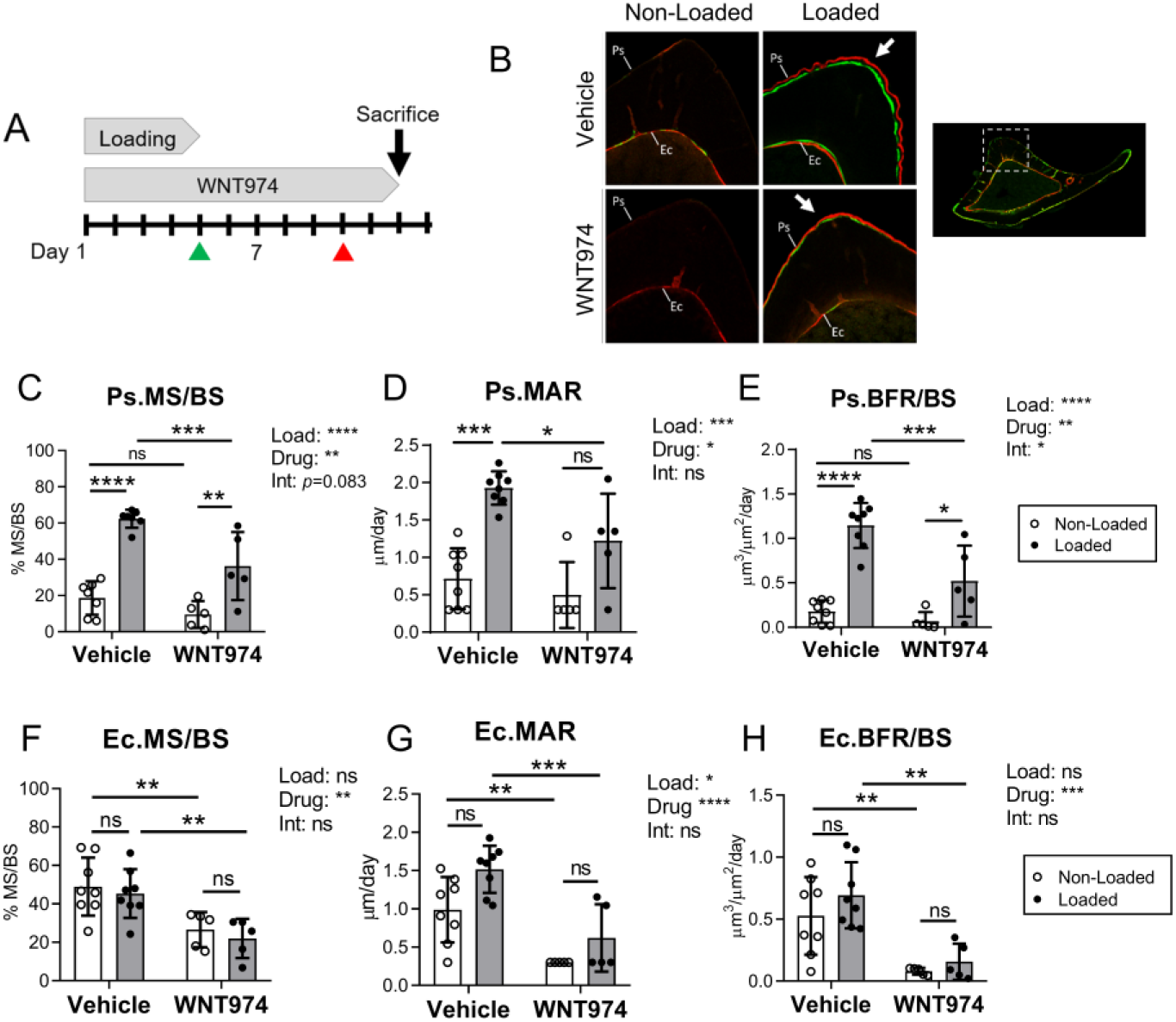
Loading-induced bone formation is diminished in WNT974-treated mice. (A) Five-month-old C57BL/6 female mice were subjected to 5 days of tibial loading, and concurrently treated with WNT974, to systemically inhibit Wnt secretion, or vehicle. Calcein and alizarin were given by IP injection on days 5 and 10, respectively (green and red arrowheads). (B) Representative images of transverse sections from the mid-diaphysis of non-loaded and loaded tibias in each group showing the region of peak compressive strain, where double-labeled surface is seen in the loaded bones (arrows). Ps= periosteum, Ec=endocortex. (C-E) Indices of periosteal bone formation showed that loading stimulated bone formation in both treatment groups, albeit significantly less in WNT974-treated mice. (F-H) Indices of endocortical bone formation show that tibial loading had a negligible anabolic effect, whereas WNT974 treatment had a profound inhibitory effect. Two-factor ANOVA was used to evaluate the effects of loading (“Load”) and WNT974 treatment (“Drug”), and their interactions (“Int”), with Sidak’s multiple comparisons test for pairwise comparisons. Bars depict mean ± SD, with individual data points shown (n=5-8). *p<0.05, **p<0.01, ***p<0.001, ****p<0.0001, ns=not significant (p>0.05).

On the *endocortical* surface, which experiences a lower strain magnitude than the periosteal surface, loading had a negligible effect on bone formation (Fig. 1F-H). Two-factor ANOVA indicated that loading did not affect Ec.MS/BS and Ec.BFR/BS and only marginally increased Ec.MAR. In contrast, WNT974 significantly reduced indices of bone formation, independent of loading. Ec.MS/BS, Ec.MAR, and Ec.BFR/BS were all significantly lower in the non-loaded tibias of WNT974-treated mice compared to vehicle-treated mice. Therefore, independent of loading, Wnt secretion is required for the maintenance of basal endocortical bone formation in 5-month old C57BL/6 mice.

### Loading-induced upregulation of osteogenic genes is reduced in WNT974-treated mice

To better understand the mechanisms underlying the osteo-anabolic deficits in WNT974-treated mice, tibial gene expression was assessed by qPCR after day 5 of loading. In vehicle-treated mice, loading potently stimulated expression of genes associated with collagen synthesis (*Col1a1*), osteoblast differentiation (*Sp7*/Osterix), and osteoblast activity (*Bglap*/Osteocalcin), whereas in WNT974-treated mice, loading failed to significantly upregulate expression of *Col1a1* or *Bglap,* and upregulation of *Sp7* was blunted (Fig. 2A-C). Moreover, two-factor ANOVA indicated that the expression of these genes was significantly reduced by WNT974 treatment in both non-loaded and loaded tibias. In contrast, neither loading nor WNT974 treatment affected expression of the late osteoblast/osteocyte marker *Dmp1* (Fig. 2D). In summary, consistent with periosteal bone formation results, loading induced the upregulation of osteogenic genes, while treatment with WNT974 strongly diminished the osteoinductive effects of loading.

**Figure 2.**
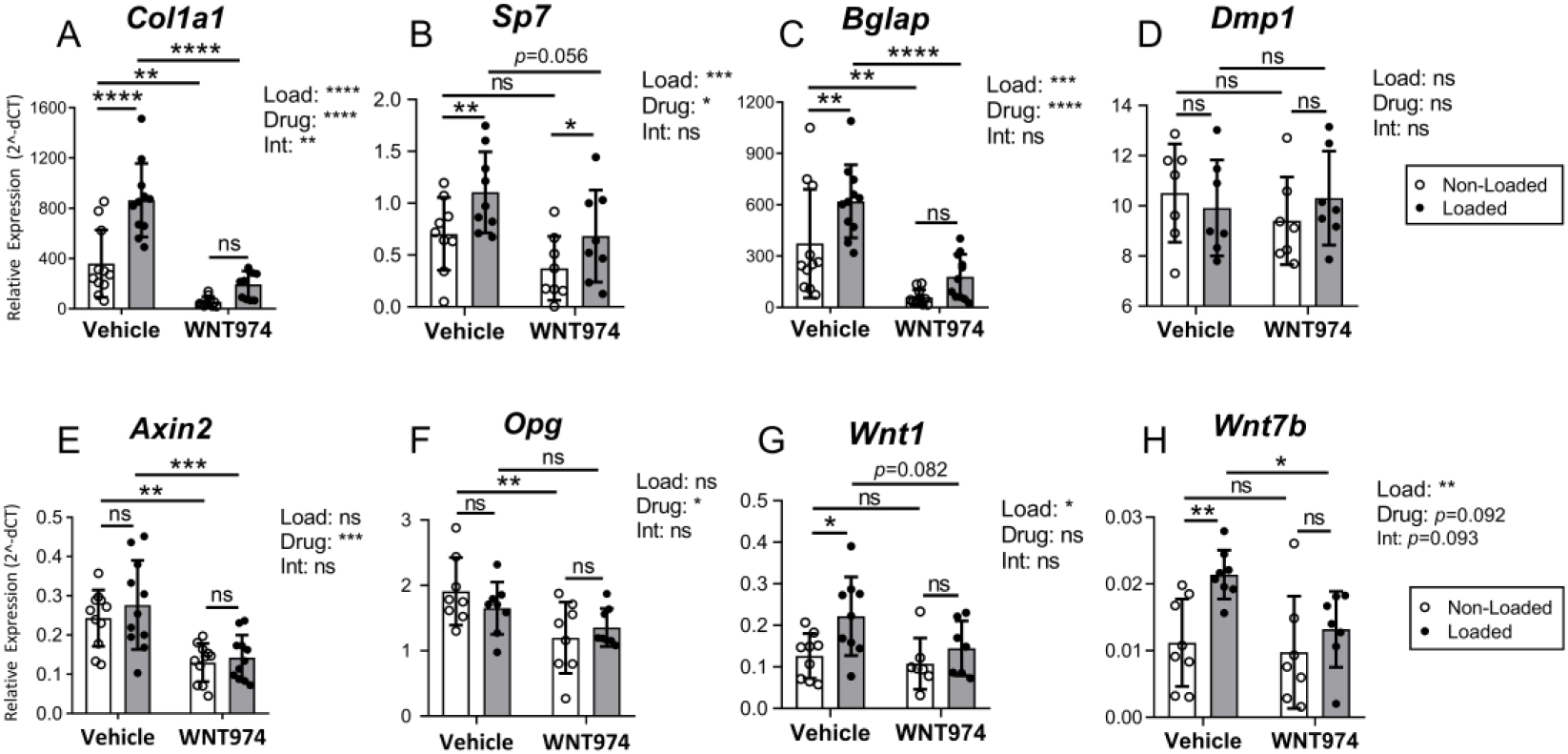
Loading-induced gene expression is altered in WNT974-treated mice. Gene expression in tibias was analyzed by qPCR to evaluate the effect of 5 days of loading and WNT974 treatment. (A-D) Tibial loading had a potent inductive effect on osteogenic gene expression. In vehicle-treated mice, *Col1a1, Sp7,* and *Bglap* were significantly higher in loaded versus non-loaded tibias. By contrast, in WNT974-treated mice tibial loading failed to significantly increase *Col1a1* and *Bglap* expression. *Dmp1* was not affected by loading in either group. (E-F) Wnt target genes *Axin2* and *Opg* were not significantly regulated by loading but were significantly lower in the bones of WNT974-treated mice (G-H). *Wnt1* and *Wnt7b* were responsive to loading but not WNT974 treatment, n=7-11/group. Data analysis as described in Figure 1. *p<0.05, **p<0.01, ***p<0.001, ****p<0.0001, ns=not significant (p>0.05).

We also surveyed a panel of Wnt pathway-related genes. Loading had a negligible effect on tibial expression of the Wnt target genes *Axin2* and *Opg,* whereas WNT974 significantly decreased their expression (Fig. 2E-F). *Axin2* and *Opg* expression were reduced by 47% and 37%, respectively, in the non-loaded tibias of WNT974-treated mice compared to vehicle-treated mice. Other Wnt pathway related genes were not consistently regulated by loading or WNT974 treatment (Suppl. Fig. S2). In contrast, two-way ANOVA indicated that *Wnt1* and *Wnt7b* were not affected by WNT974 treatment but were significantly up-regulated by loading (Fig. 2G-H). These data indicate that Wnt secretion regulates the expression of some Wnt target genes in bone, but not the expression of *Wnt1* and *Wnt7b* transcripts.

### Loading-induced bone formation is reduced in osteoblast-specific *Wls* knockout mice

To investigate the role of Wnts secreted by osteoblasts in loading-induced bone formation, we conditionally deleted the intracellular Wnt transporter Wntless *(Wls),* in Osterix-expressing cells in 5-month old female and male mice. A fluorescent Cre reporter bred into a subset of tamoxifen-inducible OsxCreERT2;Wls^F/F^ mice showed robust Cre activity in the tibia following a 3-day regimen of tamoxifen (Suppl. Fig. S3). PCR of genomic DNA showed that Wls deletion was specific to bone (Suppl. Fig. S4). Based on these results, a 3-day tamoxifen dosing regimen was used to conditionally delete Wls in adult mice, followed by loading 3 days later (Fig. 3A). Body weight did not differ between control and Wls knockouts (not shown). *Wls* mRNA expression was 53% lower in knockout tibias relative to controls at the start of loading (*p*<0.001, Fig. 3B), and likewise immunostaining for Wls protein showed reduced expression in bones of knockout mice (Suppl. Fig. S5). In addition, western blot analysis showed reduced phosphorylated-Lrp6 in knockout bones, confirming diminished activation of Wnt/Lrp signaling (Suppl. Fig. S6).

**Figure 3.**
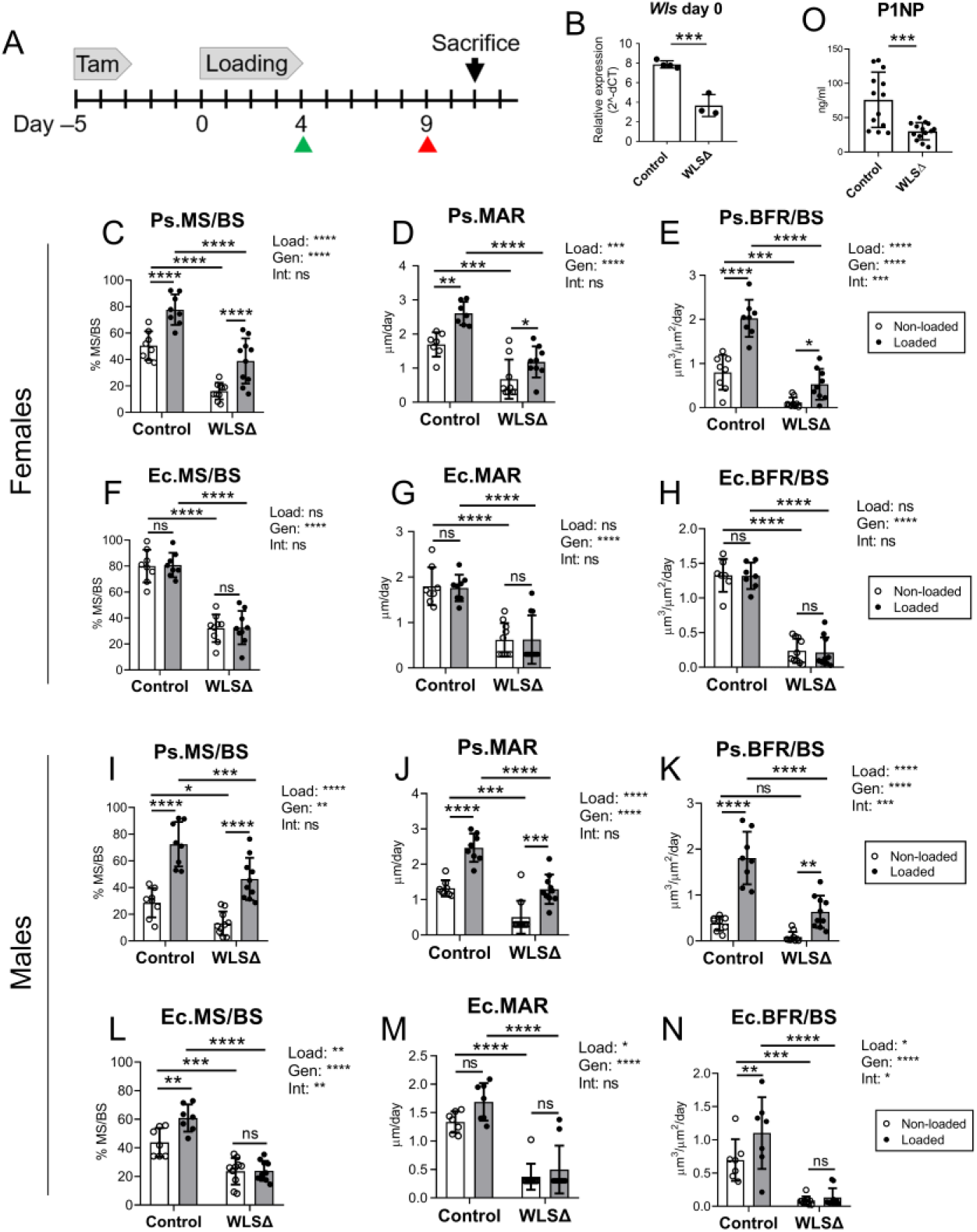
Loading-induced bone formation was impaired in Wntless knockout mice. (A) Male and female mice were given tamoxifen to induce Wls knockout at 5-months old, subjected to 5 days of tibial loading, then given calcein and alizarin (green and red arrowheads) by IP injection. (B) Tibial *Wls* RNA expression was 53% lower in Wls cKO mice (WLSΔ) compared to controls (n=3-4). (C-E) Periosteal (Ps) and (F-H) endocortical (Ec) bone formation indices in female mice (n=7-10). (I-K) Periosteal and (L-N) endocortical bone formation indices in male mice (n=8-11). (O) Serum P1NP was 60% lower in Wls cKO mice compared to controls (n=13-14). Two-factor ANOVA was used to evaluate the effects of loading (“Load”) and genotype (“Gen”), and loading-genotype interactions (“Int”), with Sidak’s multiple comparisons test for pairwise comparisons. Bars depict mean ± SD, with individual data points shown (n=5-8 per treatment group), *p<0.05, **p<0.01, ***p<0.001, ****p<0.0001, ns=not significant (p>0.05).

Next, we subjected control and Wls cKO mice to tibial loading at −3500με peak compressive strain to stimulate lamellar bone formation (Suppl. Fig. S7). In both loaded and non-loaded bones, periosteal and endocortical bone formation indices were significantly lower in *Wls* knockouts. In females, Ps.BFR/BS and Ec.BFR/BS were 85% and 82% lower, respectively, in the non-loaded tibias of knockouts relative to controls (Fig. 3E, 3H). Similarly, in males Ps.BFR/BS and Ec.BFR/BS were 78% and 87% lower, respectively, in the non-loaded bones of knockouts relative to controls (Fig. 3K, 3N). Serum analysis by ELISA showed that P1NP was 60% lower in Wls cKO mice compared to controls (*p*<0.001, Fig. 3O). Together, these data support a role for osteoblast-derived Wnts in maintaining basal-level bone formation in the adult skeleton.

Loading significantly enhanced *periosteal* bone formation in both control and Wls cKO mice, but bone formation indices were significantly lower in the loaded bones of knockout mice compared to controls (Fig. 3C-E and I-K). In females, Ps.MS/BS was 50% lower in loaded knockout bones relative to controls, while Ps.MAR and Ps.BFR/BS were 55% and 74% lower, respectively (*p*<0.0001, Fig. 3C-E). Similarly, in males Ps.MS/BS, Ps.MAR, and Ps.BFR/BS were 36%, 48%, and 65% lower, respectively, in loaded bones of knockouts vs controls (*p*<0.001, Fig. 3I-K). Additionally, a highly significant loading-genotype interaction was detected for Ps.BFR/BS in both sexes, indicating that the effect of loading on periosteal bone formation depended on genotype (*p*<0.001, Figs. 3E, 3K). This effect was further illustrated by a significant reduction in the relative (loaded minus non-loaded) periosteal bone formation rate in Wls knockouts compared to controls (Suppl. Fig. S8).

In contrast to the highly anabolic effect of loading on the periosteal surface, loading had a negligible effect on *endocortical* bone formation in females (Fig. 3F-H), and a modest effect in males (Fig. 3L-N). In males, loading significantly increased Ec.MS/BS and Ec.BFR/BS in control but not knockout mice, and two-factor ANOVA indicated a significant loading-genotype interaction for Ec.MS/BS and Ec.BFR/BS (Figs. 3L, 3N). Taken together with the periosteal results, these data demonstrate that osteoblast-derived Wnts contribute to loading-induced bone formation.

Recent studies have demonstrated a role for periosteal cell proliferation in the anabolic response to loading (Zannit and Silva, 2019) (Zannit, Brodt, et al 2020). To test whether the impaired loading response in Wls cKO mice was due to diminished cell proliferation, we dosed mice with EdU throughout the 5-day loading period and for 3 additional days until sacrifice (Suppl. Fig. S9A). Loading caused an increase in the EdU+ periosteal surface, to a similar level (~30%) in control and Wls knockout mice (Suppl. Fig. S9C). Thus, loading-induced periosteal cell proliferation appeared normal in Wls cKO mice. Similarly, we did not observe any evidence of increased TUNEL+ staining in Wls knockout bones, suggesting that their impaired anabolism was not due to increased apoptosis (Suppl. Fig. S10).

### Loading-induced expression of osteogenic and Wnt target genes is impaired in osteoblast-specific Wls knockout mice

To further characterize the effects of Wls deletion in osteoblasts, expression of genes related to bone formation and Wnt signaling was surveyed by qPCR. Wls deletion had a significant effect on osteoblastic gene expression in non-loaded bones, where *Colla1, Sp7, Bglap,* and *Pdpn* were 35-55% lower in Wls cKO mice relative to controls (*p*<0.05, Fig. 4B-E). Tibial loading potently induced osteogenic gene expression in control mice. *Bmp2, Colla1, Sp7, Bglap, Pdpn,* and *Dmp1* were all significantly higher in the loaded vs. non-loaded bones of control mice (Fig. 4A-F). In stark contrast, loading failed to significantly upregulate these same genes in Wls knockouts. A significant loading-genotype interaction was detected for *Bmp2*, *Col1a1, Sp7, Bglap,* and *Pdpn* (Fig. 4A-E), indicating that osteoblast-specific Wls expression was required for loading-induced upregulation of these osteogenic factors.

**Figure 4.**
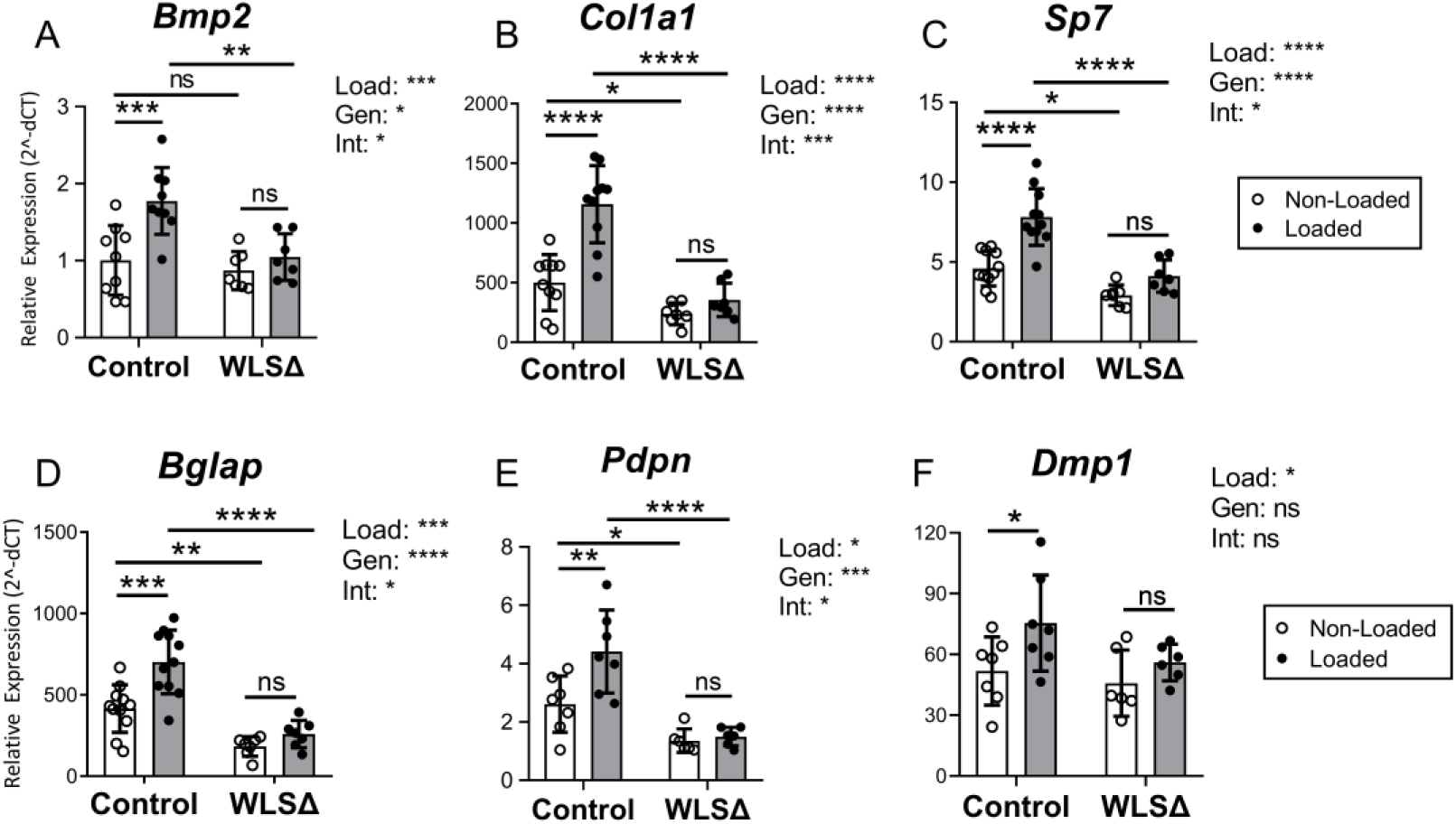
Loading-induced upregulation of osteogenic genes is impaired in Wntless knockout mice. qPCR was used to evaluate gene expression after 5 days of loading in control and Wls cKO (WLS△) mice. Expression of osteogenic-related genes was significantly higher in the loaded versus non-loaded bones in control mice, whereas loading failed to significantly upregulate osteogenic gene expression in the tibias of knockout mice. Two-factor ANOVA identified a significant loading-genotype interaction for *Bmp2, Colla1, Sp7, Bglap,* and *Pdpn,* indicating that osteoblast-specific *Wls* expression was required for the regulation of these genes by loading. Data analysis as described in Figure 3. (n=8-11) *p<0.05, **p<0.01, ***p<0.001, ****p<0.0001, ns=not significant (p>0.05).

A survey of several Wnt target genes showed that Wls deletion blunted their expression in non-loaded and loaded bones. In non-loaded tibias, *Axin2* expression was marginally lower in knockout versus control mice (−32%, *p*=0.083; Fig. 5A), while *Lrp5* and *Nkd2* were significantly lower (−32% and −60%, respectively, *p*<0.0001; Fig. 5B-C). Notably, expression of these three genes was significantly higher in loaded bones of control mice, but was not elevated in loaded bones of knockouts (Fig. 5A-C). In partial contrast, *Ccnd1* expression was increased by loading in both genotypes, although its expression was reduced in the loaded bones of knockout compared to control mice (Fig. 5D). Thus, Wls deletion in osteoblasts reduced expression of Wnt target genes in bone and their induction by mechanical loading.

**Figure 5.**
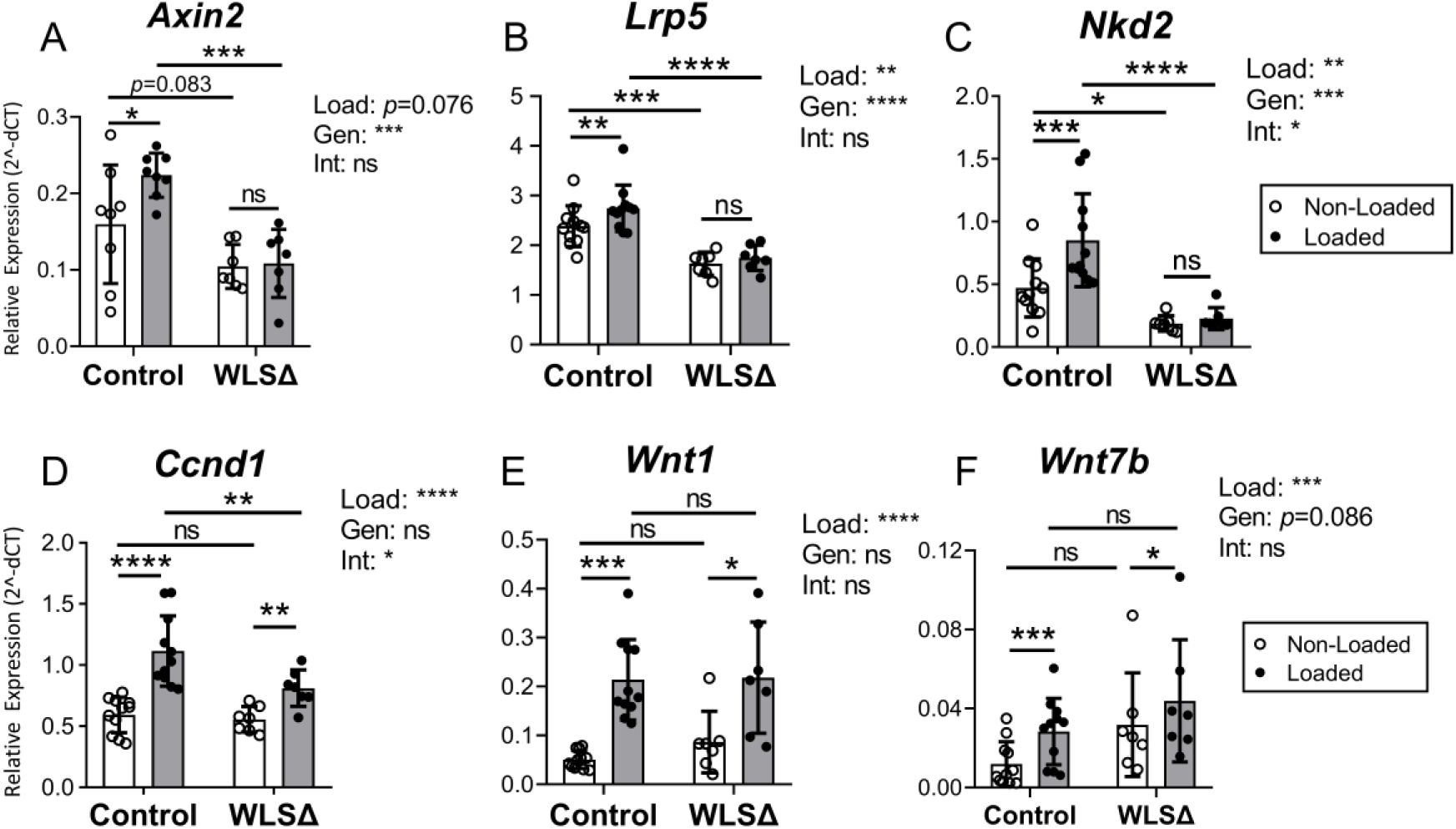
Wnt target gene expression was blunted in Wls knockouts. Expression of Wnt target genes (A-D) and Wnt pathway-related genes (E-F) in the bone was evaluated by qPCR after 5 days of loading. Wnt target genes *Axin2, Lrp5,* and *Nkd2* were significantly upregulated by loading in control but not Wls cKO (WLSΔ) mice (AC). A significant loading-genotype interaction for *Nkd2* and *Ccndl* was detected by two-factor ANOVA, suggesting a role for osteoblast-derived Wnts in regulating the expression of these genes (C-D). In contrast, *Wnt1* and *Wnt7b* were significantly higher in the loaded versus non-loaded tibias of both groups (n=8-11). Data analysis as described in Figure 3. *p<0.05, **p<0.01, ***p<0.001, ****p<0.0001, ns=not significant (p>0.05).

Recent studies report that *Wnt1* and *Wnt7b* are osteogenic (Luther, Yorgan et al. 2018) (Chen et al, 2014; Song et al. 2020), and that loading potently stimulates their expression (Kelly, et al, 2016; Chermside-Scabbo et al, 2020). Consistent with these latter reports, in control mice *Wnt1* expression was 4.2-fold higher in the loaded versus non-loaded limb, while *Wnt7b* was 2.6-fold higher (Fig. 5E-F). In knockout mice, *Wnt1* and *Wnt7b* also increased significantly with loading (2.5 and 1.4-fold higher, respectively), and there was no significant loading-genotype interaction. Together, these data confirm that loading potently induces *Wnt1* and *Wnt7b*, and demonstrate that loading-associated regulation of *Wnt1* and *Wnt7b* in the bone does not require Wnt protein secretion by osteoblasts.

Lastly, we examined the expression of several Wnt antagonists to check whether they were modulated by Wls deletion. Expression of *Sost, Dkk1, Dkk2* and *sFrp1* were either not different or were reduced in tibias of Wls cKO mice compared to controls (Suppl. Fig. S11). In addition, we stained tibial sections for Sclerostin, and observed no effect of genotype on the number of Sclerostin-positive osteocytes (Suppl. Fig. S12). Expression of these Wnt antagonists was not modulated by loading in Wls cKO mice. Therefore, the impaired bone formation in Wls knockout mice does not appear to be due to altered expression of Wnt antagonists.

### Wls deletion in osteoblast lineage cells enhances bone resorption in adult mice

Wnts produced by osteoblasts modulate bone resorption indirectly by regulating the expression of the osteoclast inhibitor *Opg,* and directly by regulating the activity of osteoclast progenitors (Moverare-Skrtic et al, 2012). In mice, constitutive Wls deletion in Ocn-expressing cells increases markers of bone resorption (Zhong, et al., 2012). To assess the short-term impact of Wls deletion in osteoblasts in adult animals, qPCR was used to analyze expression of several osteoclast regulators in the tibias of control and knockout mice. Additionally, a separate group of naïve 5-month old mice were tamoxifen-treated and sacrificed 1 week later for serum analysis.

*Opg* expression in bones of Wls knockout cKO was marginally decreased (−25%, *p*=0.052; Fig. 6A). Loading had a modest but significant stimulatory effect on *Opg* expression in bones of control but not knockout mice (*p*<0.05 versus *p*=0.46, Fig. 6A). In addition, Wls deletion increased expression of the pro-osteoclast factor *Rankl* in bone (Fig. 6B). Notably, the *Rankl* / *Opg* ratio was 2-fold higher in Wls knockout bones relative to controls (*p*<0.001), indicating that loss of Wnt secretion by osteoblasts favors osteoclastogenesis.

**Figure 6.**
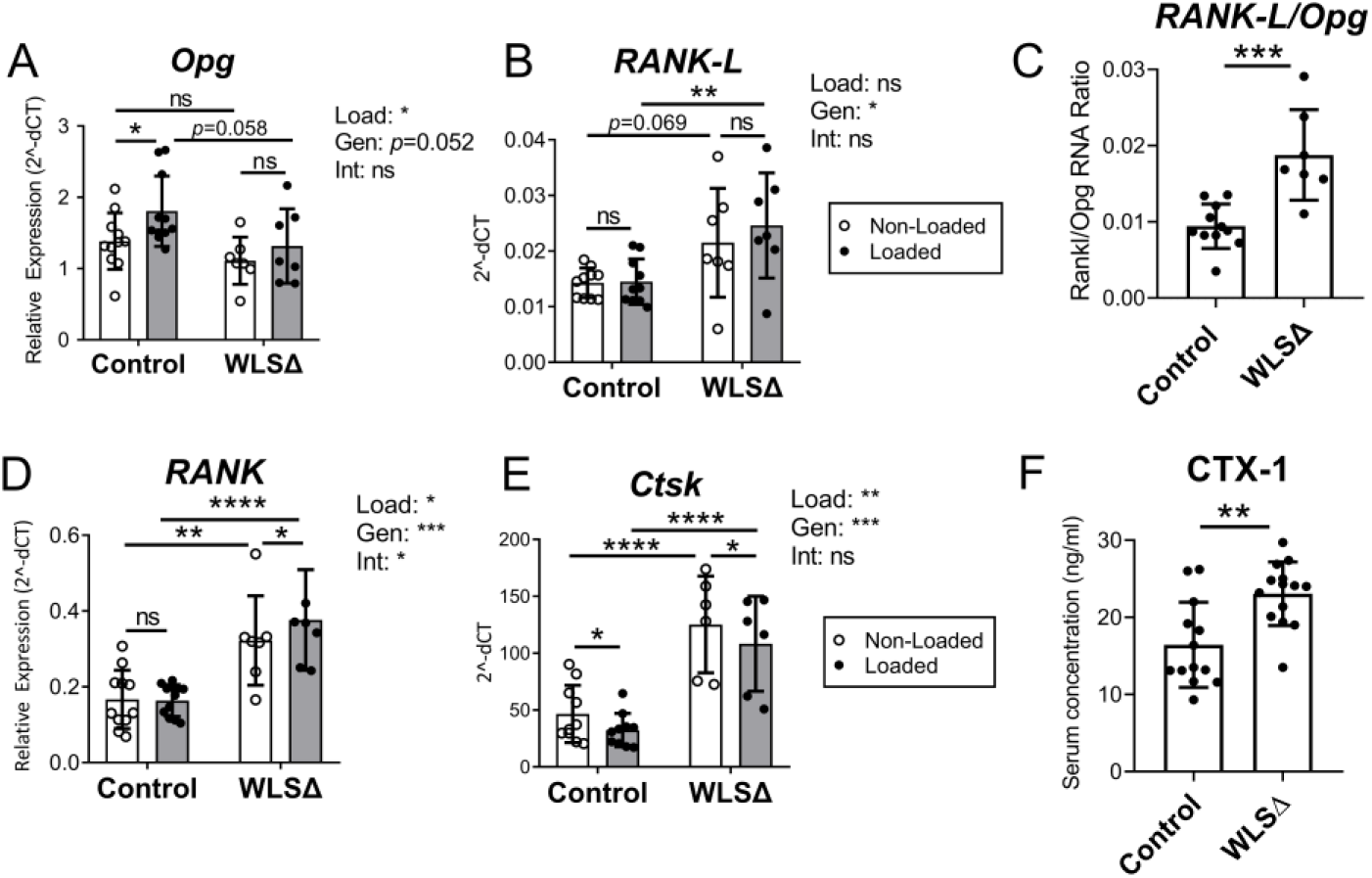
Wls deletion in osteoblasts increased bone resorption in adult mice. (A-E) Genes expressed by osteoblasts *(Opg, Rankl)* and osteoclasts *(RANK, Ctsk)* were analyzed by qPCR (n=6-10), and two-factor ANOVA was used to evaluate the effects of loading (“Load”) and genotype (“Gen”) on gene expression. Independent of loading, two-factor ANOVA indicated that loading had a modest but significant role in regulating *Opg, RANK,* and *Ctsk* in the bone, while genotype was a significant factor regulating *RANKL, RANK,* and *Ctsk. (C)* The ratio between *RANK-L* RNA and *Opg* RNA was used as a measure of local bone resorption dynamics. In non-loaded tibias, the *RANK-L* to *Opg* ratio was significantly higher in Wls cKO (WLSΔ) bones, indicating a shift toward increased bone resorption in Wls knockout mice. (F) Serum collected from naïve (not loaded) mice showed that Wls deletion significantly increased the bone resorption marker CTX-1 one week after knockout (n=13-14). Data analysis as described in Figure 3. *p<0.05, **p<0.01, ***p<0.001, ****p<0.0001, ns=not significant (p>0.05).

Expression of *RANK* and *Ctsk,* which are predominantly expressed by osteoclast lineage cells, was also elevated in Wls cKO bones. *RANK* expression was 93% greater in the non-loaded bones of knockouts versus controls (*p*<0.01), while *Ctsk* was 260% higher (*p*<0.0001) (Fig. 6D-E). Finally, serum CTX-1 was 40% higher in naïve (non-loaded) Wls cKO mice compared to controls (*p*<0.01, Fig. 6F). Together these data indicate that even brief interruption of Wnt secretion in osteoblasts has a significant impact on bone turnover in the adult skeleton.

## Discussion

Wnt signaling is critical to many aspects of skeletal regulation, but the importance of Wnt ligands in adult bone homeostasis and the anabolic response to mechanical loading is not well documented. We inhibited Wnt ligand secretion in adult (5-mo) mice using a systemic (drug) and a bone-targeted (genetic) approach, and subjected them to axial tibial loading. Mice treated with the porcupine inhibitor WNT974 exhibited a decrease in bone formation in non-loaded limbs as well as a marked decline in the anabolic response to tibial loading. Similarly, within 1-2 weeks of Wls deletion in osteoblasts, skeletal homeostasis was altered with decreased bone formation and increased resorption, and the anabolic response to loading was greatly impaired. These findings establish a requirement for Wnt ligand secretion by osteoblasts for adult bone homeostasis and the anabolic response to mechanical loading.

Previous reports have established a requirement for Wnt ligand secretion in skeletal development and maturation. Treatment of 8-12 wk old mice using the porcupine inhibitor WNT974 reduces bone mass within 3-4 weeks (Funck-Brentano et al., 2018; Madan et al., 2018). Furthermore, Wls deletion in osteoblasts severely impairs skeletal development leading to perinatal lethality in Osx-Cre;Wls cKO mice (Tan et al., 2014), and severe osteopenia and skeletal fragility within 2 months postnatally in Ocn-Cre;Wls cKO mice (Zhong et al., 2012). Our results extend these findings to adult (5-mo) mice. Porcupine inhibition led to reduced bone formation (independent of mechanical loading), evident by a significantly lower expression of osteogenic genes (e.g, *Col1a1, Sp7)* and rates of endocortical bone formation in the non-loaded limbs of WNT974 treated mice. Furthermore, when we deleted Wls in osteoblasts of 5-mo old mice using the inducible Osx-CreERT2 driver, within 1-2 weeks we observed reduced osteogenic gene expression in bone, a lower serum level of the bone formation marker P1NP, an increase in the *Rankl/Opg* ratio in bone and in the serum marker of bone resorption CTX-1, along with reduced periosteal and endocortical bone formation rates in non-loaded bones of Wls cKO mice. Collectively, these results establish that secretion of Wnt ligands by osteoblasts is essential for skeletal homeostasis in adult mice, and that short-term inhibition of Wnt secretion causes rapid, deleterious changes in bone turnover.

The ability of the bone to adapt to mechanical stimuli is a key principal of skeletal physiology. A main objective of this study was to assess the requirement of Wnt ligand secretion for the anabolic response to mechanical loading. The importance of the Wnt signaling pathway in bone’s loading response has been demonstrated in LRP5 mutant mice (Niziolek et al., 2012; Robinson et al., 2006; Sawakami et al., 2006; L. Zhao et al., 2013), and with the finding that overexpression of the Wnt antagonist SOST (normally downregulated by loading (Robling et al., 2008)) blocks the anabolic loading response (Tu et al., 2012). Several studies show that loading upregulates the expression of Wnt genes including *Wnt1, Wnt7b, Wnt10b* and *Wnt16* (Chermside-Scabbo et al., 2020; Galea et al., 2017; Holguin et al., 2016; Kelly et al., 2016; Robinson et al., 2006; Wergedal et al., 2015), suggesting a functional role for increased ligand abundance in the bone formation response. Notably, Wergedal et al. (Wergedal et al., 2015) reported that 10-wk old mice with global deletion of *Wnt16* had a diminished periosteal response to tibial bending (Wergedal et al., 2015). Here, we extend these studies to show that Wnt ligand secretion is essential for the anabolic response to axial tibial loading in adult mice. First, we observed an increase in osteogenic gene expression after 5 days of loading in control mice, which is a normal precedent to increased bone formation (Chermside-Scabbo et al., 2020; Holguin et al., 2016; Mantila Roosa et al., 2011). But this loading-induced increase in osteogenic gene expression was diminished in both WNT974 treated and Wls cKO mice. For example, loading increased the expression of type I collagen transcript *Col1a1* by 2.3-fold in control mice, but this effect was absent in Wls cKO mice. Moreover, tibial loading significantly increased *periosteal* bone formation rate in control mice, an effect which was significantly diminished in both WNT974 treated and Wls cKO mice. Loading also increased *endocortical* bone formation in male control mice but not in Wls cKO mice. Taken together, these results demonstrate a requirement for Wnt ligand secretion by osteoblasts in the anabolic response to loading. Moreover, these results motivate follow-up studies targeting osteoblast expression of individual! Wnt ligands to identify which ones are important in the bone anabolic response.

Results from Wls cKO mice offer insights into the regulation of Wnt related gene expression by Wnt signaling in osteoblasts. Non-loaded bones from Wls cKO mice had reduced expression of the canonical Wnt target gene *Axin2* and also showed a trend for lower levels of phosphorylated LRP6, consistent with reduced Wnt signaling (Clevers & Nusse, 2012). Wls cKO led to reduced expression of Wnt pathway antagonists *Dkk1, Sost* and *Nkd2* (Burgers & Williams, 2013; Canalis, 2013; S. Zhao et al., 2015), suggesting that Wnt signaling positively regulates these genes as part of a feedback loop. Notably, expression of *Wnt1* and *Wnt7b* was not significantly affected by Wls deletion. Moreover, these ligands were upregulated by loading in both control mice (consistent with previous reports (Chermside-Scabbo et al., 2020; Holguin et al., 2016; Kelly et al., 2016)), and also in Wls cKO mice. This finding suggests that load-induced upregulation of these two ligands is not regulated by Wnt signaling in osteoblasts. Additional work is needed to clarify the mechanisms whereby Wnt ligands in bone are induced by mechanical loading, although one recent study found that the upregulation of Wnt1 in osteocytes was regulated by the mechanosensitive Piezo1 channel, possibly via YAP/TAZ (Li et al., 2019).

Our findings highlight the functional importance of Wnt ligand secretion in skeletal regulation. Recent genome-wide association study (GWAS) results identified 89 variants in the human WLS gene as being significantly associated (either positively or negatively) with estimated bone mineral density (eBMD) in the UK Biobank dataset (Morris et al., 2019; “Musculoskeletal Knowledge Portal,”). In contrast, there were no variants in PORCN that were significantly associated with eBMD. The association of WLS variants with bone mass in humans, in combination with the current and previous loss of function studies in mice (Zhong, et al. 2012; Tan et al., 2014), suggests that minor dysregulation in Wnt ligand secretion may influence bone mass accrual in response to anabolic cues, including mechanical loading.

While responses to loading were diminished by targeting Wnt secretion, they were not absent. Periosteal bone formation indices were significantly higher in loaded vs. non-loaded limbs of WNT974 treated as well as Wls cKO mice, a result that might be due to two factors. First, non-Wnt pathways are also important in loading-induced bone formation, as noted in Sost and Dkk1 knockout mice (Pflanz,et al., 2017; Morse et al., 2020), and second, Wnt secretion was not completely blocked in our studies. We cannot distinguish the relative importance of these two by factors, but note that even a ~50% reduction in *Wls* expression in bone of cKO mice (Fig. 3B) is sufficient to reduce their relative loading-induced periosteal bone formation rate by 60-70% (Suppl. Fig. S8).

In summary, we used pharmacological (WNT974) and genetic (Wls cKO) approaches in adult mice to disrupt Porcupine and Wntless, two factors required for Wnt ligand secretion. Mice were subjected to unilateral axial tibial loading, an established model for induction of bone formation. Local bone responses were assessed by qPCR and dynamic histomorphometry, while systemic responses were assessed by serum markers of bone turnover. Results from WNT974 treated mice and Wls cKO mice were consistent, showing sharply diminished measures of bone formation in both non-loaded and loaded bones compared to control mice. In particular, the normal induction of bone formation by mechanical loading seen in control mice was significantly blunted in mice with inhibition of Wnt ligand secretion. On the other hand, the loading-induced upregulation of *Wnt1* and *Wnt7b* were not impaired by the interventions. We conclude that secretion of Wnt ligands by osteoblasts is required for homeostasis and the full anabolic response to mechanical loading in the adult skeleton.

## Methods

### Animal models

Studies were approved by the Washington University IACUC. Mice were housed up to five per cage and given ad libitum access to normal chow and water. Mice were euthanized by CO2 asphyxiation.

#### Porcupine inhibition

A pharmacological approach was used to inhibit Wnt secretion in 5-month old C57BL/6 female mice (Charles River) assigned randomly to two groups. Porcupine inhibitor WNT974 (aka Lgk974; Active Biochemicals, Wan Chai, Hong Kong) was prepared as previously described (Zhang and Lawrence, 2016) and delivered daily by oral gavage at a dose of 6mg/kg/day. Control mice received an equivalent volume of vehicle (0.15ml, 5% carboxymethylcellulose (w/v) in water).

#### Osteoblast-specific Wls deletion

An inducible Cre/LoxP approach was used to conditionally delete *Wls* in Osterix-expressing cells. OsxCreERT2 sires (derived from breeders gifted from Dr. Henry Kronenberg) were mated to dams that were homozygous for the conditional floxed *Wls* allele (JAX #012888) to generate OsxCreERT2; Wls^F/F^ experimental (Wls cKO/WLSΔ) and Wls^F/F^ control littermates. Mice were genotyped by Transnetyx (Transnetyx, Inc. Cordova, TN, USA) using tail snip DNA with probes for wildtype *Wls (Gpr177-1 WT),* conditional *Wls* (*Gpr177-1 FL*), and OsxCreERT2 (*Cre*). After genotyping, mice were assigned to cKO or control groups as available until the desired sample sizes for each outcome were attained (see below), with similar numbers of littermates of each genotype included in each experimental batch. Tamoxifen was dissolved in corn oil to a final concentration of 10mg/ml and was delivered by oral gavage at a dose of 50mg/kg/day for 3 consecutive days to induce gene knockout in 5-month old male and female mice. Tamoxifen-treated Wls^F/F^ mice served as genotype controls for all experiments. The inducible Ai9 reporter transgene (JAX #007909) was introduced into a subset of mice to generate OsxCreERT2; Wls^F/F^; Ai9 mice, which were used to assess Cre specificity. All mouse strains have been previously described (Carpenter et al., 2013; Maes et al., 2007; Madisen et al., 2010).

### Tibial loading

In vivo tibial compression was used to stimulate cortical bone formation in 5-month old mice, using a 5-day loading protocol as previously described (Sun et al., 2019; Main et al., 2020). Briefly, an Electropulse 1000 system (Instron, USA) was used to apply 60 cycles (4Hz) of axial compression on the right tibias of anesthetized mice (mouse prone/tibia vertical). Our goal was to engender an estimated −2200μe peak strain at the 37% diaphyseal cross-section (5 mm proximal to the distal tibiofibular junction) to induce lamellar bone formation in C57BL/6 mice (Sun et al., 2019; Holguin et al., 2014). Based on *a priori* strain gauge measurements (Suppl Fig. S7) the peak forces to accomplish strain-matched loading were: −8N WLSΔ and control females, and −11N WLSΔ and control males. The mice were subjected to tibial loading for 5 consecutive days. Contralateral left tibias served as non-loaded controls. Mice designated for gene expression analysis were sacrificed 4 h after the final bout of loading. Mice designated for dynamic histomorphometry received calcein (10mg/kg; Sigma) by IP injection on the last day of loading, followed by alizarin complexone (30mg/kg; Sigma) 5 days later.

### Fluorescent histology

#### Dynamic histomorphometry

Bilateral tibias were plastic-embedded as described (Brodt and Silva 2010) and sectioned transversely at the 37% location to a thickness of 100 □m. Images at a resolution of 2048×2048 were captured on a Leica confocal microscope (see below).

Fluorophore-labeled surfaces were analyzed using commercial software (Osteo II, Bioquant) to calculate standard indices of bone formation, including percent mineralizing surface (MS/BS), mineral apposition rate (MAR), and bone formation rate (BFR/BS) (Dempster et al,. 2013). Values were determined on total periosteal (Ps) and endocortical (Ec) surfaces separately. In instances where no double-labeled surface could be detected, or where the inter-label distance was too small to measure, a value of 0.3μm/day was assigned (Dempster et al., 2013). In addition to absolute values for loaded and non-loaded tibias, relative bone formation rate (BFR_Loaded_ – BFR_Non-Loaded_) was calculated as a measure of the anabolic response to loading. For all quantitative histomorphometry, the user was blinded to group assignment prior to analysis.

#### Cre reporter analysis

Ten □m thick transverse sections from the non-loaded tibias of tamoxifen-treated OsxCreERT2; Rosa^Ai9/+^; Wls^F/F^ mice were DAPI-counterstained and imaged on a Leica confocal microscope to analyze endogenous Ai9 reporter expression 3 days after the final dose of tamoxifen.

#### EdU and TUNEL analyses

A commercially available kit (Click-iT EdU Cell Proliferation Kit, ThermoFisher) was used to label proliferating cells with thymidine analog 5-ethynyl-2’- deoxyuridine (EdU). Briefly, EdU (0.2mg/ml) dissolved in 5% sucrose water (w/v) was delivered daily to the mice via drinking water during the loading period until sacrifice. Tibias harvested 3 days after loading were fixed in 4% paraformaldehyde (PFA), de-calcified in 14% EDTA, incubated in 30% sucrose/PBS overnight, and embedded in OCT (optimal cutting temperature) medium. Twenty □m thick sections sectioned on a cryostat were stained for EdU following manufacturer instructions. Sections were DAPI-counterstained and imaged on a confocal microscope. Percent proliferation was defined as the ratio between EdU-positive nuclei and total (DAPI-positive) cells on the bone surface. Cell death was analyzed similarly using a commercially available kit that labels fragmented DNA from apoptotic cells (DeadEnd Fluorometric TUNEL System, Promega).

All fluorescence imaging was done on a Leica inverted laser scanning confocal microscope (Leica DMi8/TCS SPE) using a 10μm Z-stack.

### Gene expression

#### Sample preparation

Gene expression was analyzed by qPCR as described (Holguin et al., 2016). Briefly, mice were sacrificed on the last day of loading, 4 h after tibial compression. Bilateral tibias were dissected out and carefully stripped of muscle, and cut 2mm distal to the tibial plateau and at the tibiofibular junction (TFJ). Bones were microcentrifuged to remove bone marrow, then placed in liquid nitrogen. Samples were pulverized with a Mikro Dismembrator (Braun) and lysed in TRIzol. Total RNA was extracted and purified using the Qiagen RNeasy Mini Kit. Complementary DNA (cDNA) was prepared with the iScript cDNA Synthesis Kit (BioRad).

#### RT-qPCR

Gene expression was analyzed by Sybr- and Taqman-based qPCR. Sybr-based qPCR was run on a StepOne Plus Machine (Applied Biosystems). Taqman-based qPCR was run on a BioMark HD System in collaboration with the Genome Technology Access Center (GTAC) at Washington University in St. Louis. Genes were chosen to assess osteogenesis *(Bmp2, Col1a1, Sp7, Bglap, Pdpn* (E11), *Dmp1),* osteoclast activity *(Opg, Rankl, Rank, Ctsk)* and Wnt signaling *(Axin2, Ccnd1, Ctnnb1, Dkk1, Dkk2, Lrp5, Lrp6, Nkd2, Sfrp1, Sost, Wnt1, Wnt7b).* mRNA expression was reported as relative expression (2^−ΔCt^), normalized to *Tbp.*

### *Wls* knockdown validation

First, genomic DNA (gDNA) was extracted from the skeletal and extra-skeletal tissues (DNeasy kit, Qiagen) of 5-month old Wls mice to evaluate DNA recombination after tamoxifen induction. Primers up- and downstream of the floxed locus in the *Wls* transgene (exon 1) were used to amplify wildtype (1625bp) and mutant (140bp) Wls DNA. P1:

CTTCCCTGCTTCTTTAAGCGTC; P4: CTCAGAACTCCCTTCTTGAAGC. PCR conditions were as described previously (Carpenter, Rao et al. 2010). Second, tamoxifen-treated mice were sacrificed 3 days after tamoxifen induction, and tibias were harvested and processed for qPCR as described under *Gene Expression.* Primers were used to amplify *Wls* exon 1, which is excised upon Cre-induced recombination. Thus, expression levels reflect the abundance of wild-type *Wls* RNA in the bone. Expression was normalized to *Tbp*. Primer sequences were: WLS-F: CAAATCGTTGCCTTTCTGGTG; WLS-R: TTGTCACACTTGTTAGGTCCC; TBP-F:

CTGAATAGGCTGTGGAGTAAGTC; TBP-R: CTGAAGAAAGGGAGAATCATGGA (Carpenter, et al., 2010).

## Immunohistochemistry

### Tissue preparation

Standard procedures were used to prepare paraffin-embedded tissue sections for immunohistochemistry. Briefly, bones were fixed in 4% PFA overnight at 4°C, followed by de-calcification for 7-10 days in 14% EDTA. Paraffin-embedded tissues were sectioned to a thickness of 10 μm on a microtome and mounted onto glass slides using a flotation water bath system.

### Immunostaining

Heat-induced epitope retrieval in sodium citrate buffer was used for antigen retrieval. Blocking buffer comprised of 3% BSA in PBST was used to block tissues for 1 h at room temperature. Primary antibodies, which were diluted 1:1000 in blocking buffer, included anti-Gpr177 *(Wls, Evi)* antibody RLAB-177 (Seven Hill Bioreagents) and anti-Sclerostin antibody AF1589 (R&D Systems). Tissues were incubated in primary antibody overnight at 4°C. Secondary reagents included VectaStain Elite ABC HRP kits PK-6101 and PK-6105, and ImmPact DAB peroxidase (HRP) substrate kit SK-4105 (Vector Labs). Tissues were counterstained with hematoxylin.

## Serum markers

Mice were tamoxifen-treated for 3 consecutive days, then sacrificed 1 week later for serum harvest. Commercially available ELISA kits from IDA Immunologistics were used to assay serum P1NP and CTX-1. Two-factor ANOVA indicated that sex was not a significant variable, and consequently, female and male data were pooled.

## Western blot

Bilateral tibias were collected from control and *Wls* knockout mice 3 days after tamoxifen induction; mice were not subjected to loading. Tibias were harvested as described above (in *Gene Expression*), then flushed with ice-cold PBS to remove residual marrow. Protein was extracted from bone and the abundance of phosphorylated LRP6 (pLRP6, CST #2568) and total LRP6 (tLRP6, CST #3395) were assessed by western blot. The ratio of pLRP6/tLRP6 was used as an indicator of Wnt signaling. (See Suppl. Fig. S6.)

## Data Analysis

Group sample sizes (n) were determined based on *a priori* power analysis and are indicated in the Results. Two-factor analysis of variance (ANOVA) was used to study the main effects of loading (non-loaded vs. loaded), and treatment (vehicle vs. WNT974) or genotype (control vs. WLSΔ), and their interaction, on bone formation and gene expression outcomes with Sidak’s multiple comparisons test for pairwise comparisons (Prism, Graphpad 7.0). For bone formation outcomes in the Wls deletion study, male and female mice were analyzed separately. One-factor ANOVA was used for outcomes where loading was not a factor (e.g., serum markers). Significance was defined at p<0.05. Individual data points and the mean ± standard deviation are plotted.

## Acknowledgments

This work was supported by NIH grants R01 AR047867, T32 AR060719 and the Washington University Musculoskeletal Research Center (P30 AR074992). We thank Crystal Idleburg and Samantha Coleman for histology support. We thank the Genome Technology Access Center (supported by P30 CA91842 and UL1 TR002345) for help with gene expression assays. We thank the Core Laboratory for Clinical Studies for their help with serum assays. Finally, we thank Roberto Civitelli for advice and comments on the study.

Authors’ roles: Study design: LYL, MDB and MJS. Study conduct and data collection: LYL, MDB, NM, CCS, and RP. Data analysis and interpretation: LYL, MDB, NM, CCS, and MJS. Drafting manuscript: LYL and MJS. Revising manuscript content: LYL, NM, CCS, and MJS. Approving final version of manuscript: all authors. LYL and MJS take responsibility for the integrity of the data analysis.

## Supplemental material

**Supplemental Figure S1.**
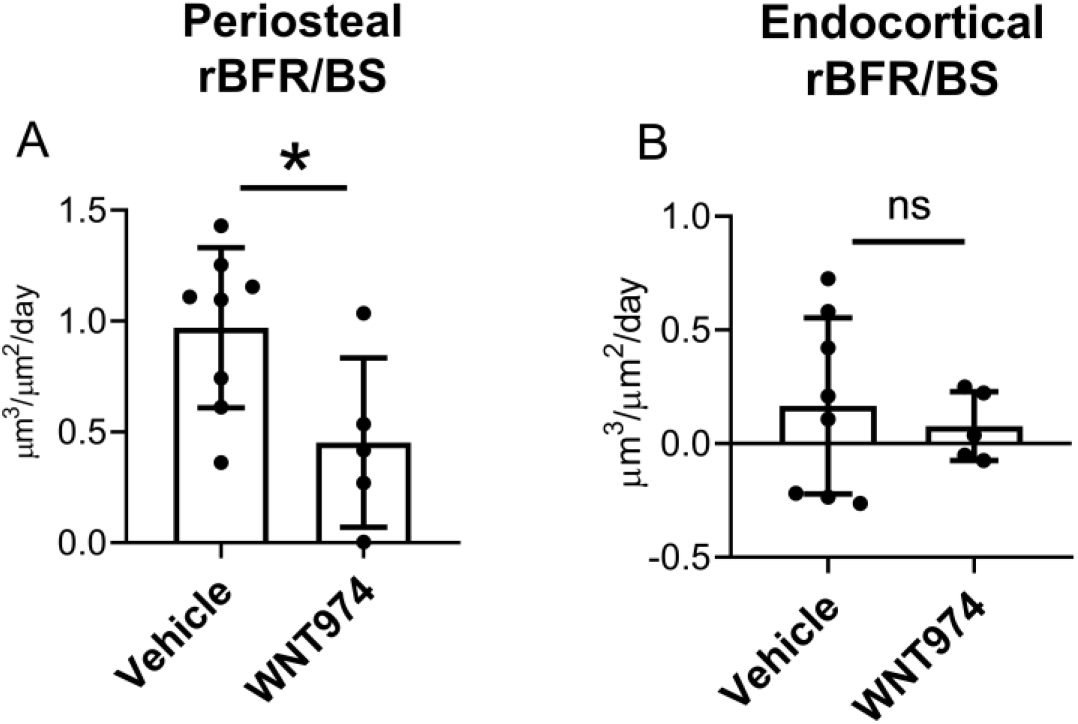
Relative bone formation rate (BFR/BS_Loaded_ minus BFR/BS_Non_Loaded_) was used as an index of loading-induced bone formation. (A) Loading-induced periosteal bone formation was reduced in WNT974-treated mice relative to vehicle-treated mice. (B) Loading had a negligible effect on endocortical bone formation in both groups. Bars depict mean ± SD, with individual data points shown (n=5-8). One-factor ANOVA; *p<0.05, ns=not significant (p>0.05).

**Supplemental Figure S2.**
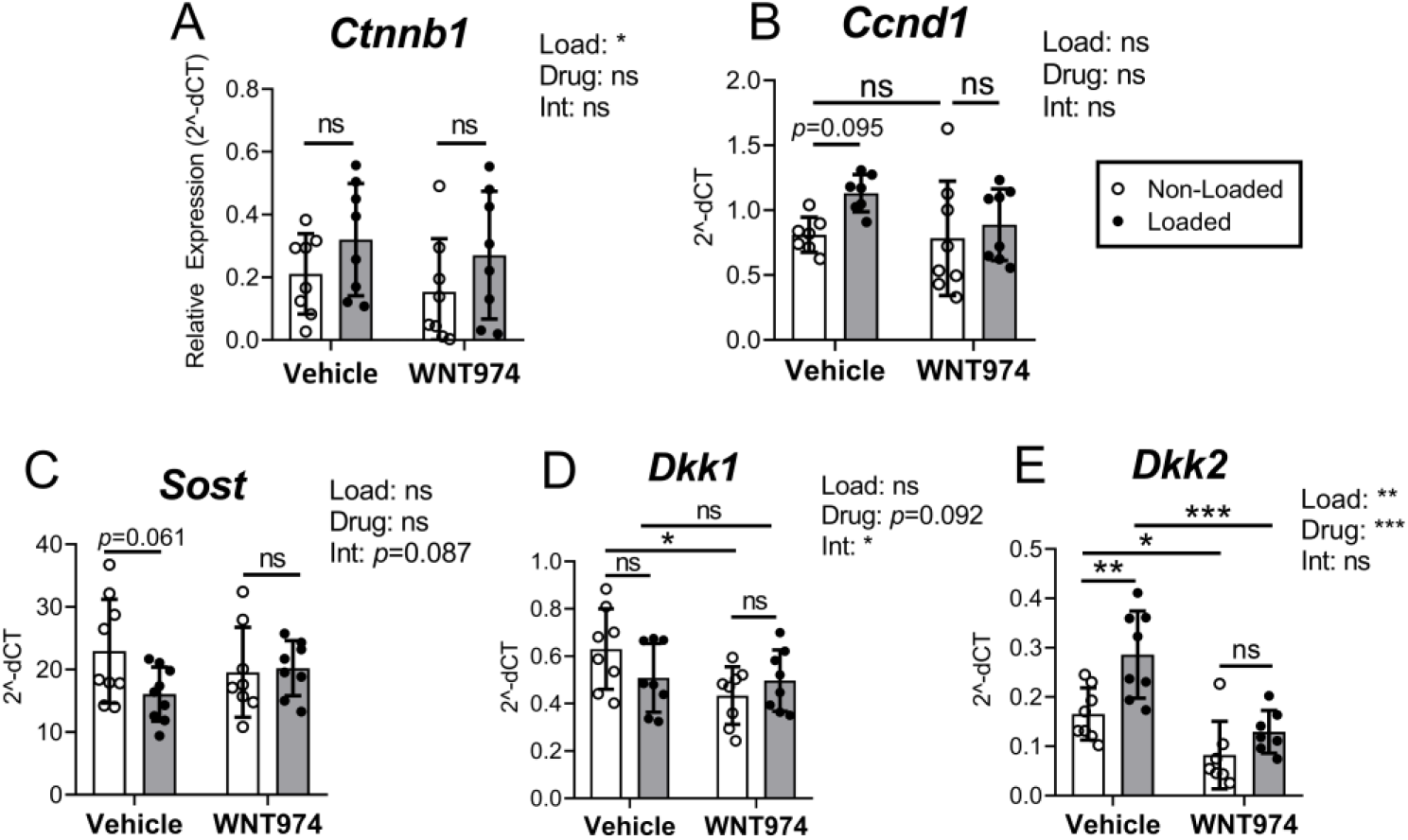
Tibial gene expression was analyzed by qPCR to evaluate the effect of 5 days of loading (“Load”), WNT974 treatment (“Drug”), and their interaction (“Int”) using two-factor ANOVA with Sidak’s multiple comparisons test. Bars depict mean ± SD, with individual data points shown (n=8-10). *p<0.05, **p<0.01, ***p<0.001, ****p<0.0001, ns=not significant (p>0.05). Data analysis as described in Figure 3.

**Supplemental Figure S3.**
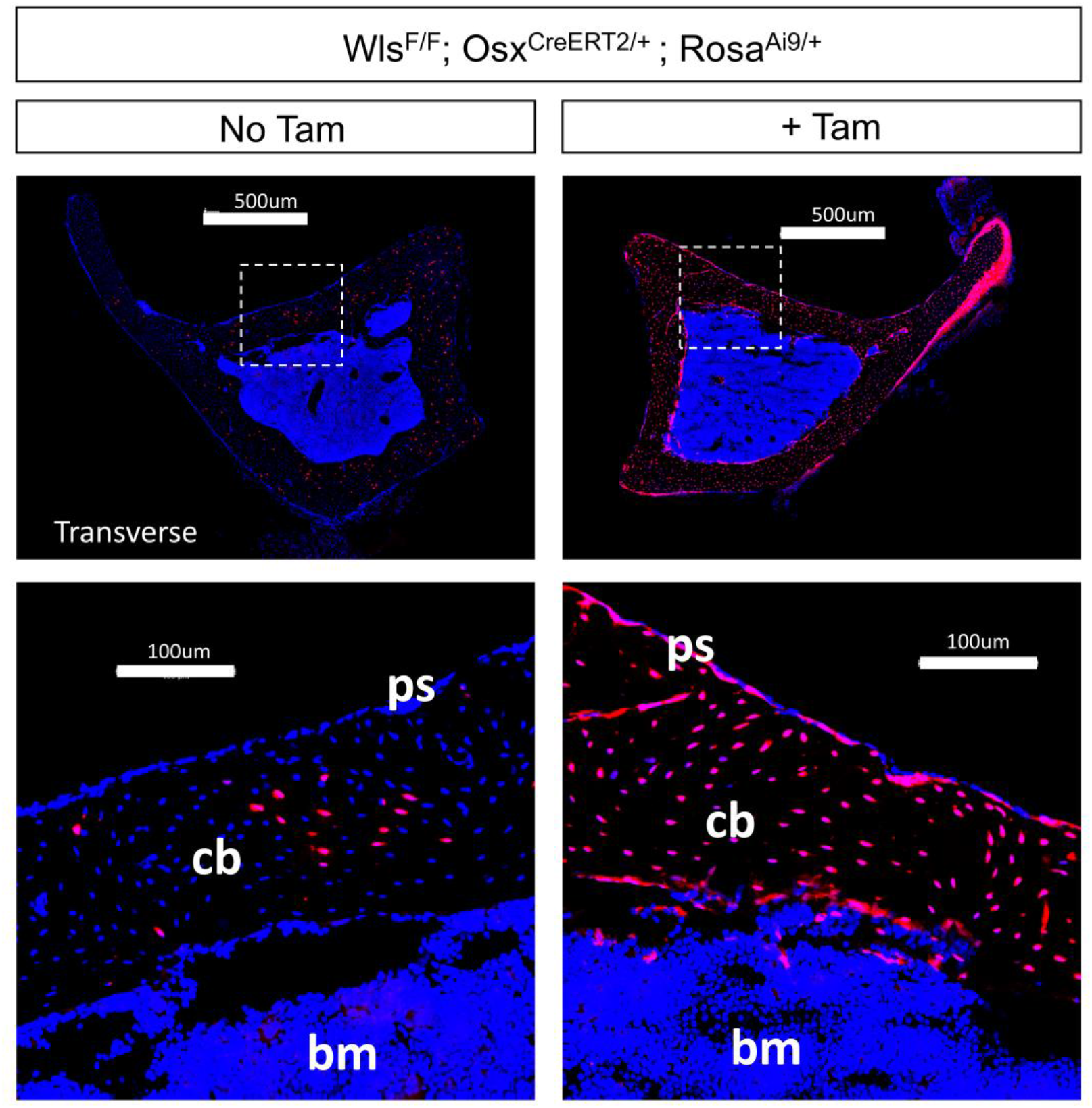
Cre reporter expression in the tibia. A fluorescent Cre-reporter was bred into the transgenic Wls colony to generate Wls^F/F^; Osx^CreERT2/+^;Rosa^Ai9/+^mice, which were treated with tamoxifen (Tam) to delete *Wls* and to survey tdTomato/Ai9 reporter fluorescence in the tibia. tdTomato expression was observed throughout the cortical bone (cb) and in the periosteum (ps) of tamoxifen-treated mice. Some tdTomato-positive cells were also observed in negative control cortical bones of mice that never received tamoxifen, ps=periosteum, cb=cortical bone, bm=bone marrow. Results are comparable to recently published results from our lab (Zannit et al., 2019).

**Supplemental Figure S4.**
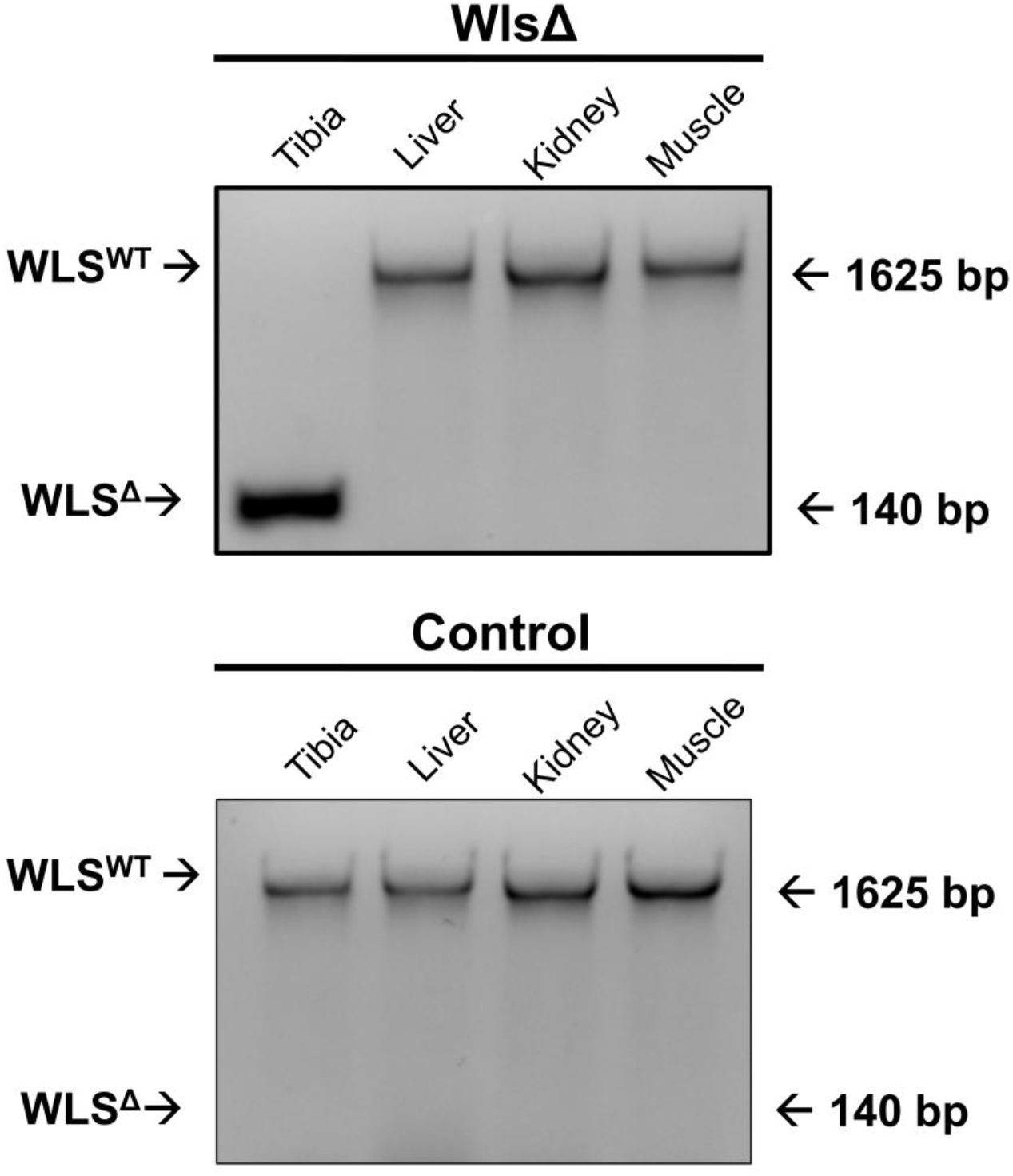
DNA recombination PCR was used to evaluate the specificity of Wls deletion. Primers up- and downstream of the floxed locus *(Wls* exon 1) were used to amplify wild-type (I625bp) and recombined (150bp) DNA. Recombination at the locus and excision of exon 1 removes the ATG start site, rendering a *Wls* null allele (WLSΔ). DNA recombination was observed in the bones of Wls knockout mice but not in the bones of control mice. No DNA recombination was observed in any of the extra-skeletal tissues surveyed from either knockout or control mice. Results are representative of n=3 Wls cKO and 1 control.

**Supplemental Figure S5.**
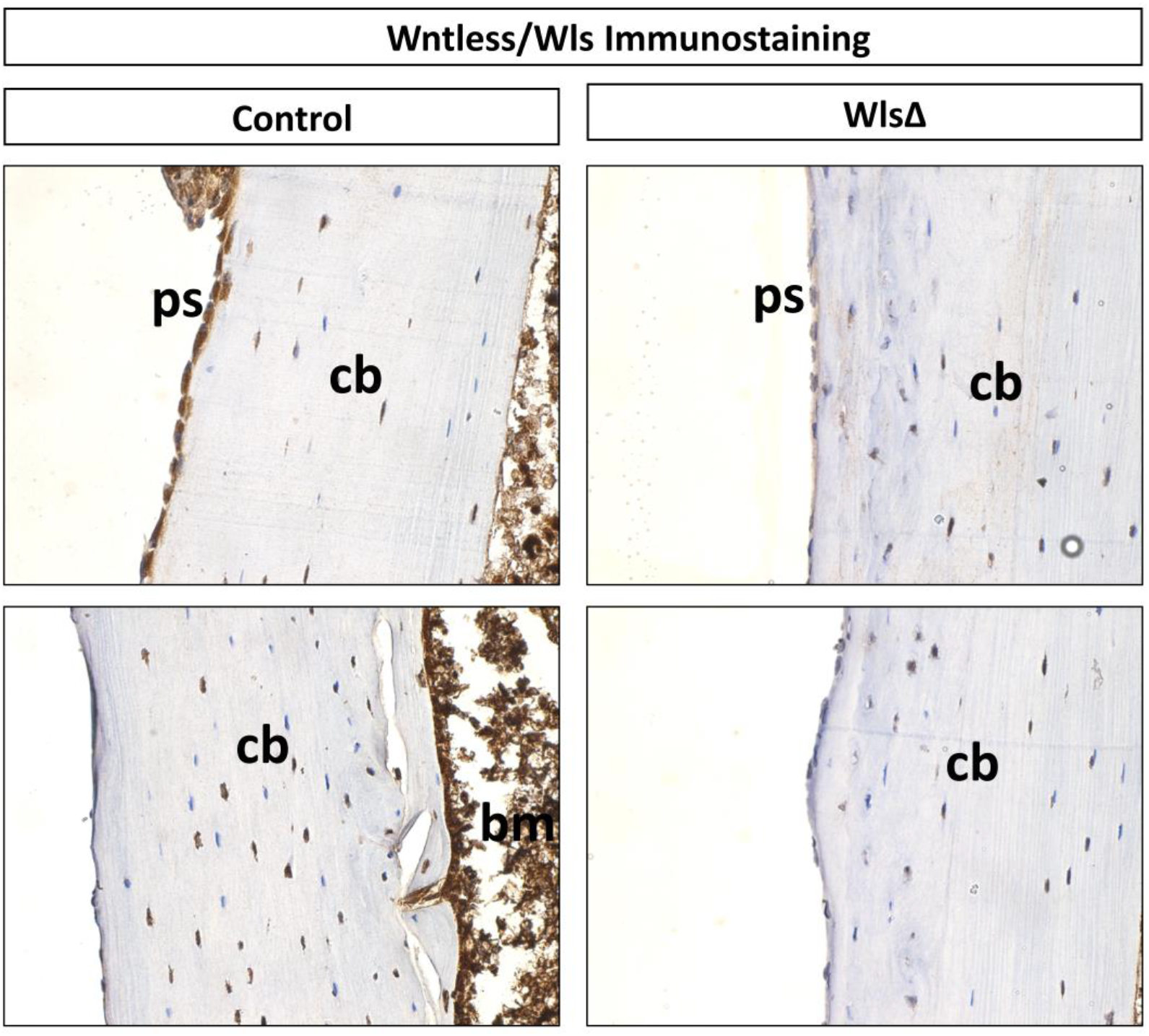
Wntless protein expression in the tibia. Sagittal sections from control and knockout mice were incubated with an antibody specific for mouse Wntless (aka GprI77) then counterstained with Hematoxylin (blue). Relative to control tissues (left two panels), there were fewer Wntless-positive cells (brown puncta) throughout the cortical bone (cb) and periosteum (ps) of *Wls* knockout bones. Tibias were sectioned through the sagittal plane and imaged at 4Ox. ps=periosteum, cb=cortical bone, bm=bone marrow. Results are representative of n=3 Wls cKO and 3 control mice.

**Supplemental Figure S6.**
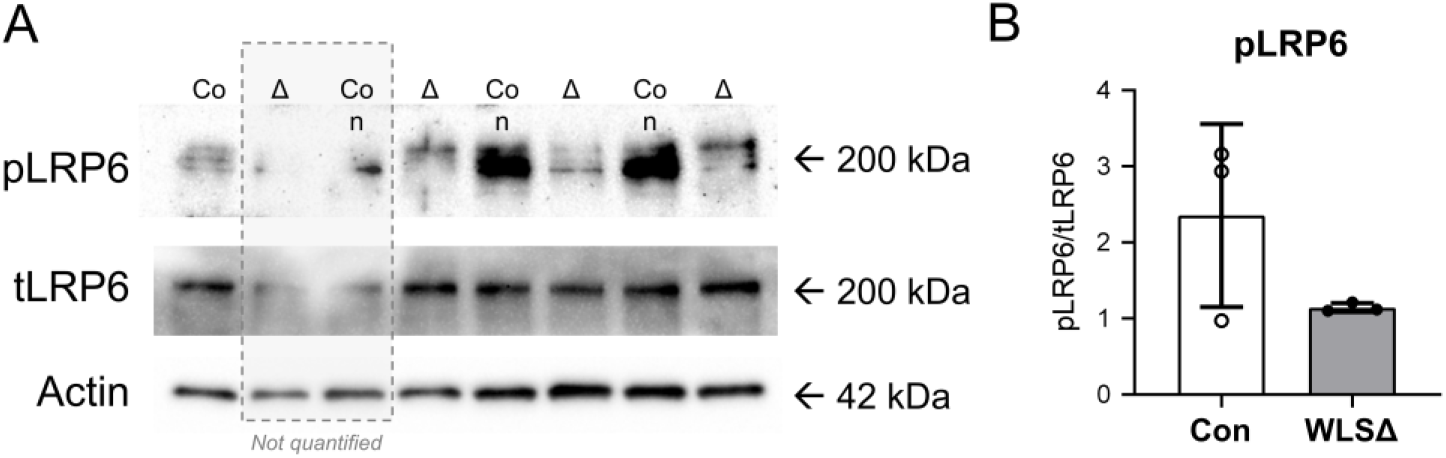
Western blotting showed diminished Wnt signaling in WLS^RF^;Osx^CreERT2^ (WLSΔ) compared to WLS^RF^ control (Con) mice. LRP5/6 are transmembrane co-receptors for Wnt ligands that are required for canonical Wnt/β-catenin signaling. Upon stimulation with Wnt ligands, LRP6 is phosphorylated at multiple sites. For this experiment, mice were gavaged with tamoxifen for 3 days, had 2 days of clearance, were sacrificed on what would have been day one of loading to assay Wnt signaling at the start of the loading. Tibias were stripped of muscle, cut at the distal tibiofibular junction (TFJ) and 2mm distal to the tibial plateau, centrifuged to remove the bone marrow, and flushed with ice-cold PBS. Protein was extracted with scissors on ice in 150μL of RIPA buffer (CST #9806) spiked with protease and phosphatase inhibitors (ThermoFisher #78440). After 25min on ice with vortexing every 5min, supernatant was collected, and ~2Oμg of protein was run on an Any kD gel (Bio-Rad #4569033). Proteins were transferred onto a PVDF membrane (wet transfer with buffer containing 20% methanol) at 4°C for 85min at constant 75V. The membrane was blocked with 2.5% BSA (Sigma #10735086001) in tris-buffered saline with 1% Tween-20 (TBS-T) for 1 hr. The membrane was cut at 75kDa and incubated with the respective primary antibodies (1:1000) for pLRP6 (CST #2568) and β-actin (CST #8457) at 4°C overnight. Membranes were washed with TBS-T (3xIOmin), incubated with HRP-conjugated anti-rabbit secondary antibody (1:2000, Sigma GENA934) in 2.5% BSA for 1 hr. After washing, ECL substrate (Bio-Rad 1705062) was used to image the membrane on the Bio-Rad ChemiDoc XRS+ platform. After acquisition, the membrane was stripped for 8min with (Theremo #46430) and probed for LRP6 (CST #3395), as described. Band intensity was quantified in ImageJ.

**Supplemental Figure S7.**
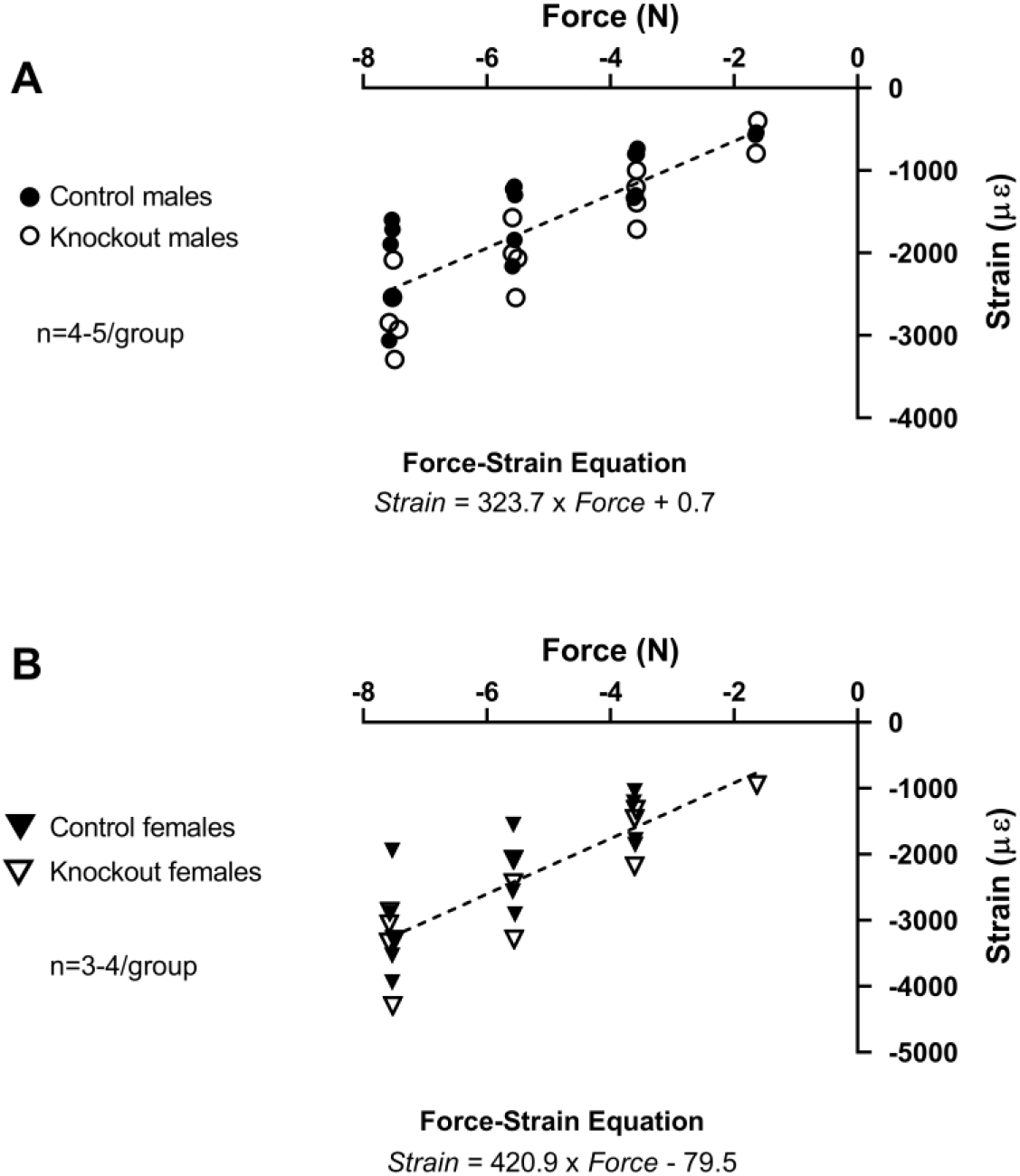
Force-strain relationships in 5-month old male (A) and female (B) Wls^RF^ control and OsxCreERT2;Wls^F/F^ cKO mice (WLSΔ). Males and females were loaded to −11N and −8N, respectively, which engendered an estimated peak compressive strain of −3500 μĉ. Regression lines were similar between control and knockout mice of each gender, and thus they were pooled for a single sex-specific regression. Males had a lower slope (stiffer) and thus required a higher force to generate an equivalent target strain compared to females.

**Supplemental Figure S8.**
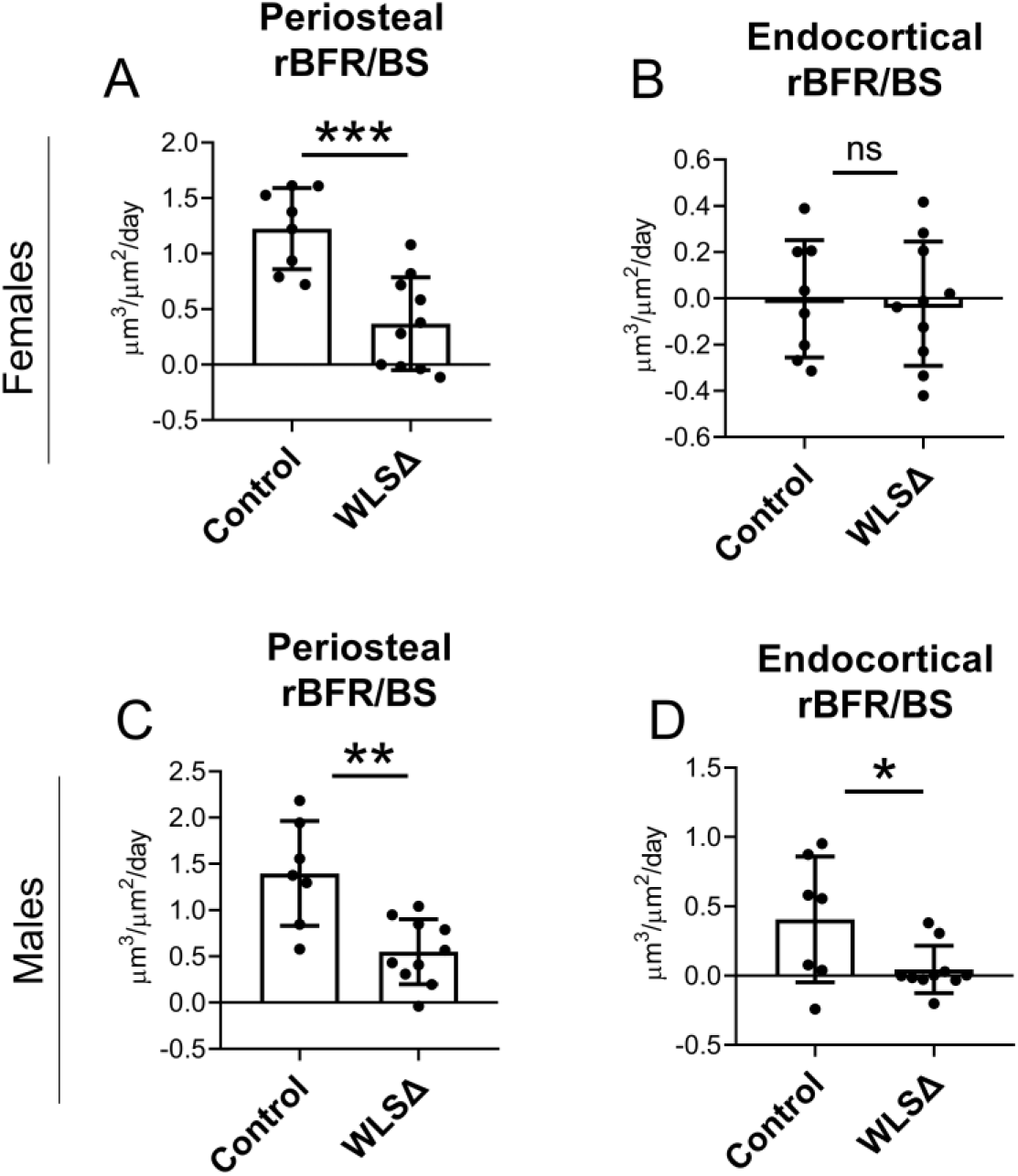
Relative bone formation rate (BFR/BS_Loaded_ minus BFR/BS_No⋂-Loaded_) was used as θ⊓ index of loading-induced bone formation. (A) In females, tibial loading stimulated periosteal bone formation in both control and knockout mice; however, loading-induced periosteal bone formation was significantly lower in knockouts (B) Tibial loading in female mice did not stimulate endocortical bone formation, regardless of genotype. (C) Loading stimulated periosteal bone formation in both control and knockout males, but rBFR/BS was significantly lower in knockouts. (D) Loading induced endocortical bone formation in control but not knockout males. Bars depict mean ± SD, with individual data points shown (n=7-10). One-factor ANOVA; *p<0.05, **p<0.01, ***p<0.001, ns=πot significant (p>0.05).

**Supplemental Figure S9.**
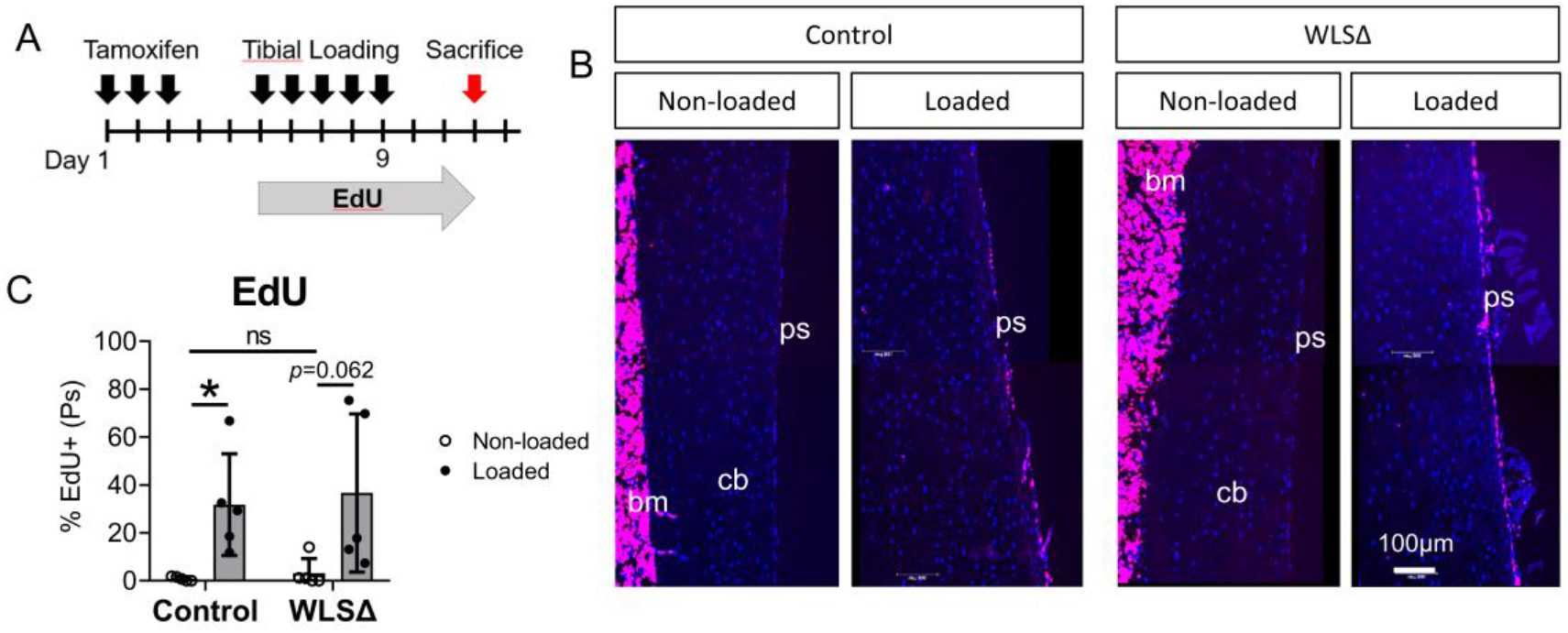
Loading-induced proliferation on the cortical bone surface was observed in both control and Wls cKO (WLSΔ) mice. (A) Mice were loaded for 5 days and EdU was administered in the drinking water to label proliferative cells. (B) Representative DAPI/EdU merged images from the non-loaded and loaded tibias of control and knockout mice shown. (C) ps=periosteum, cb=cortical bone, bm=bone marrow. Bars depict mean ± SD, with individual data points shown (n=5). One-factor ANOVA; *p<0.05.

**Supplemental Figure S10.**
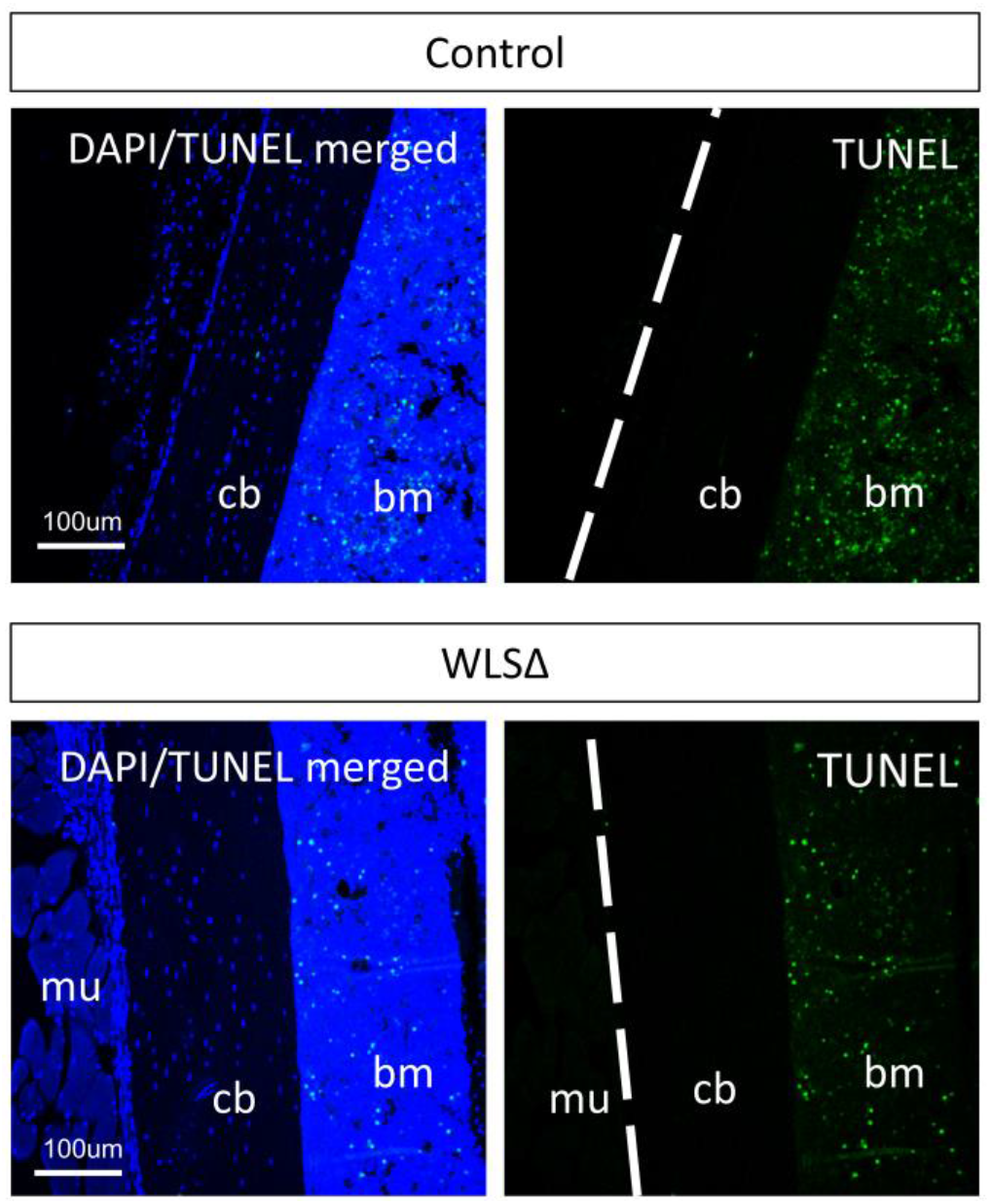
The TUNEL assay was used to assess cell death in the non-loaded and loaded tibias of control and Wls cKO (WLSΔ) mice after 5 days of loading. TUNEL staining was comparable in the non-loaded (shown) and loaded (not shown) tibias of both groups. Results are representative of n=3 non-loaded and 3 loaded tibias from each genotype, cb=cortical bone, bm=bone marrow, mu=muscle.

**Supplemental Figure S11.**
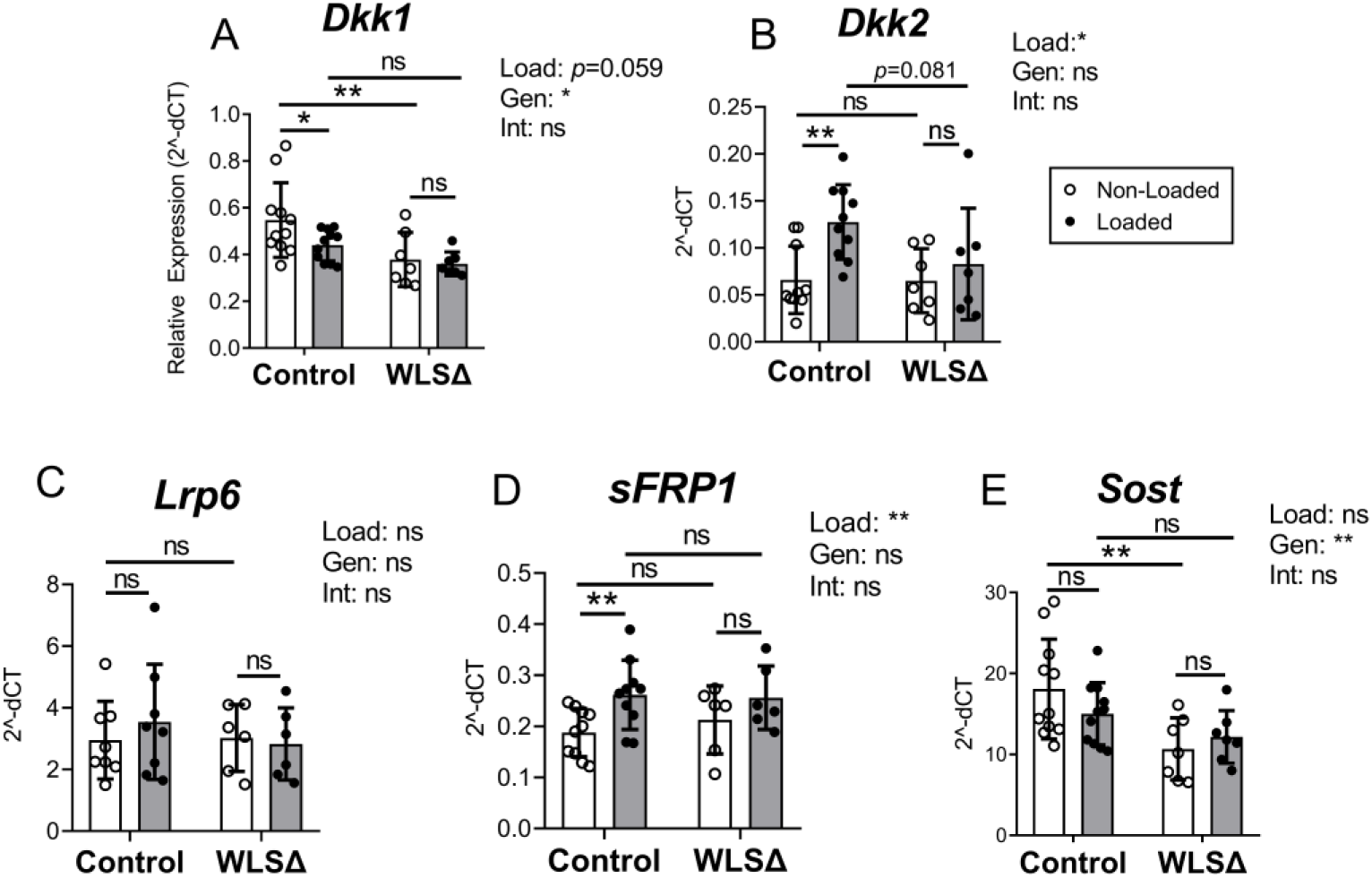
Expression of Wnt pathway inhibitors in the tibia after 5 days of loading. Gene expression was evaluated by qPCR (n=7-11). *p<0.05, **p<0.01, πs=not significant (p>0.05). Data analysis as described in Figure 3.

**Supplemental Figure S12.**
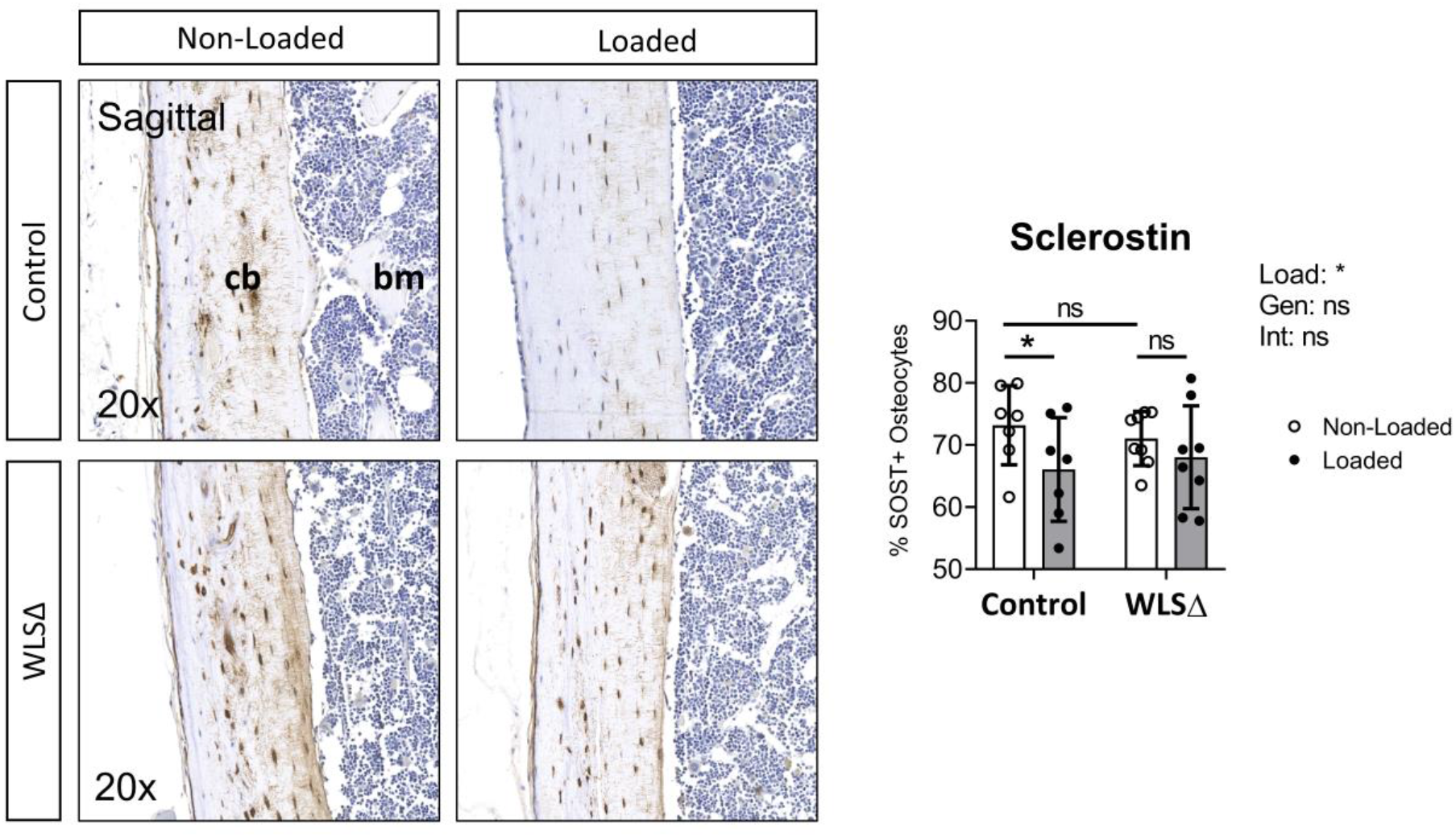
Sclerostin immunohistochemistry. An antibody specific to mouse Sclerostin was used to analyze Sclerostin protein expression in the tibias of control and WIs cKO (WLSΔ) mice. Representative images of Sost staining in the nonloaded (left) and loaded (right) tibias of control (top) and WIs cKO (bottom) mice shown (A). Percent SOST+ osteocytes was calculated as the ratio between Sost÷ osteocytes (brown puncta) to total osteocytes (blue puncta) at the mid-diaphysis (B). Bars depict mean ± SD, with individual data points shown (n=7-8) *p<0.05. Data analysis as described in Figure 3.

